# Machine Leaning-based Determination of Sampling Depth for Complex Environmental Systems: Case Study with Single-Cell Raman Spectroscopy Data in EBPR Systems

**DOI:** 10.1101/2020.12.18.423496

**Authors:** Guangyu Li, Chieh Wu, Dongqi Wang, Varun Srinivasan, David R. Kaeli, Jennifer G. Dy, April Z. Gu

**Affiliations:** Department of Civil and Environmental Engineering, Northeastern University, Boston, MA; Department of Electrical and Computer Engineering, Northeastern University, Boston, MA; Xi’an University of Technology, Xi’an, Shaanxi, PRC; School of Civil and Environmental Engineering, Cornell University, Ithaca, NY

**Keywords:** single cell technology, Single-cell Raman Microspectroscopy, Machine Learning, EBPR, Sample Size Assessment

## Abstract

Rapid progress in various advanced analytical methods such as single-cell technologies enable unprecedented and deeper understanding of microbial ecology beyond the resolution of conventional approaches. A major application challenge exists in the determination of sufficient sample size without sufficient prior knowledge of the community complexity and, the need to balance between statistical power and limited time or resources. This hinders the desired standardization and wider application of these technologies. Here, we proposed, tested and validated a computational sampling size assessment protocol taking advantage of a metric, named kernel divergence. This metric has two advantages: First, it directly compares dataset-wise distributional differences with no requirements on human intervention or prior knowledge-based pre-classification. Second, minimal assumptions in distribution and sample space are made in data processing to enhance its application domain. This enables test-verified appropriate handling of datasets with both linear and non-linear relationships. The model was then validated in a case study with eight SCRS phenotyping datasets each sampled from a different enhanced biological phosphorus removal (EBPR) activated sludge community located across North America. The model allows the determination of sufficient sampling size for any targeted or customized information capture capacity or resolution level. For example, an approximated sampling size of 50 or 100 spectra for full-scale EBPR-related ecosystems at 5% or 2% OPU cluster resolution. Promised by its flexibility and minimal restriction of input data types, the proposed method is expected to be a standardized approach for sampling size optimization, enabling more comparable and reproducible experiments and analysis on complex environmental samples. Finally, these advantages exhibit the capability of generalizing to other single-cell technologies or environmental applications, provided that the input datasets contain only continuous features.

**TOC:** 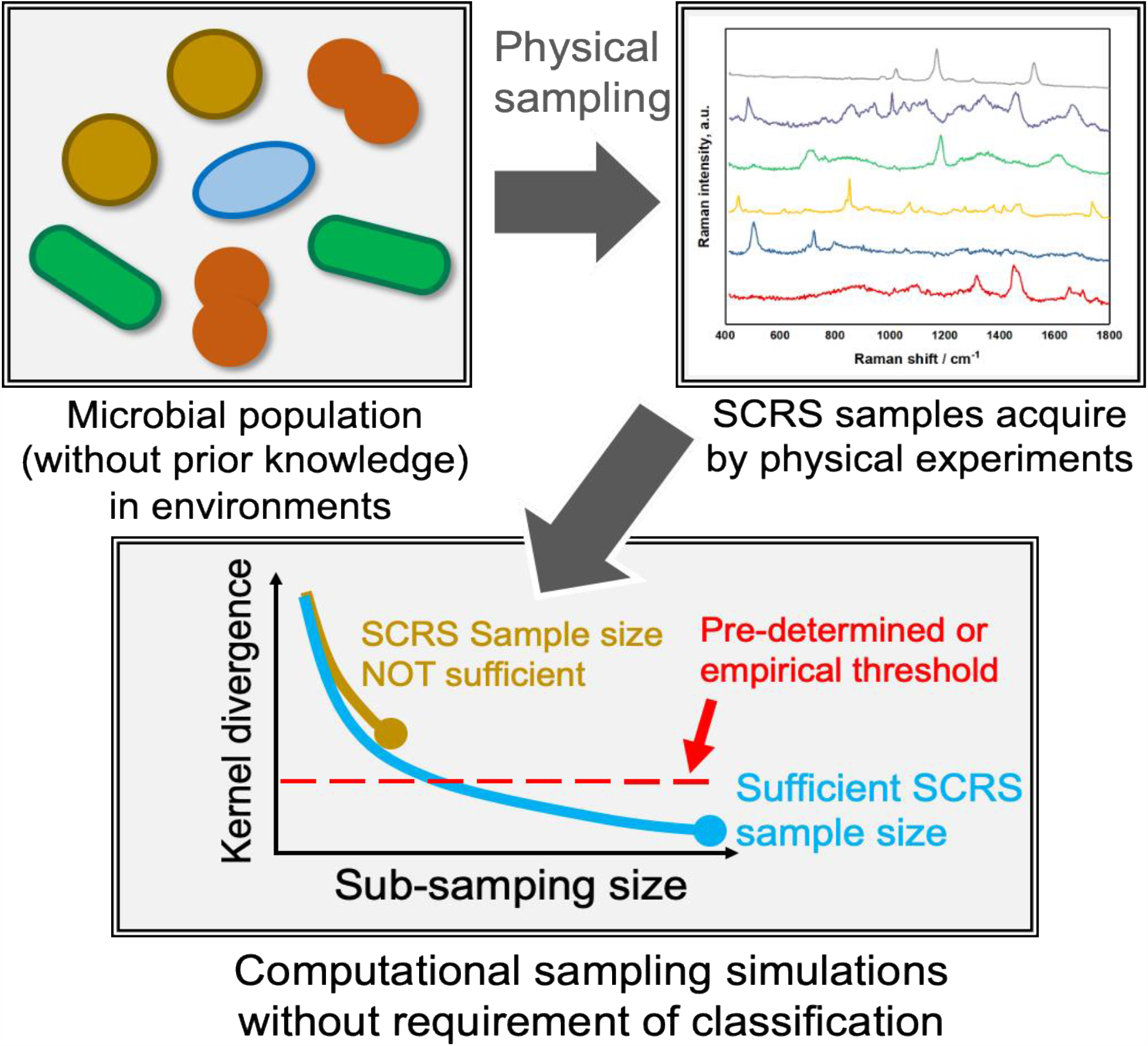

## INTRODUCTION

Advances in various modern analytical methods such as single-cell technologies have enabled unprecedented high-resolution and fundamental study of environmental microbiology than traditional cultivation-based and bulk-measurement methods. Some examples include metabolite probing (e.g. stable isotope probing) ^1, 2^, single-cell phenotype identification (e.g., fluorescence *in-situ* hybridization (FISH)) _3_ and sensitive, high through-put cell sorting (e.g., FISH-activated flow cytometry (FACS) and optical tweezer-based cell sorting) ^4, 5^. Situational studies in addition exhibit demand for non-invasive, real-time, label-free and continuously observation methods, in complement to or beyond these current single-cell technologies. These technologies, including single-cell Raman microspectroscopy can reveal cell response and metabolic changes under stimulation from various environmental changes.

Being a member of vibrational spectroscopic technology, Raman spectroscopy profiles the photons which are inelastically scattered to different frequencies due to the energy exchange between the monochromatic photon and a vibrating molecule. Its result spectra encode the fingerprints for pinpointing the chemical composition in observed cell sample, and ultimately, resolving its cellular phenotype and metabolic state. Single-cell Raman spectroscopy (SCRS) and its combination with other single-cell methods present promises for meeting this demand ^6-9^, and they have been demonstrated as powerful techniques in sub-cellular level substrate composition profiling ^10, 11^, high through-put metabolic pathway and cell type identification ^6, 12, 13^, cell sorting ^14^, and qualitative or quantitative 3D structural imaging ^15, 16^.

SCRS has been explored and demonstrated as one of the top candidate single-cell techniques, for cell identification up to strain-level discrimination of microorganism members from targeted community from different environmental matrices, including clean room ^17^, drinks ^18^, food ^19^, water ^20^, or cerebrospinal fluid ^21^. Xu and Webb et al. demonstrated that the high resolution of SCRS was able to discern two strains of only single-gene mutation apart ^22^. In addition, in contrast and complementary to genomics-based microorganism profiling approaches, SCRS captures and reflects the cell’s “metabolic state”, which is more dynamic and responsive to environmental stimuli. Furthermore, SCRS enabled technologies may help fill the gap in linking cellular phenotypes with their genotypes ^23^.

One challenge associated with the application of single-cell technology such as SCRS for complex environmental samples is the determination of sufficient sampling size without prior knowledge of the system diversity, yet with the need to balance between statistical power and cost of resources and time. This challenge raises from two major facts. First, to our knowledge, current automation level for SCRS sampling still under-satisfies the high through-put requirements for very-large scale surveys. Restricted by its labor and time demand, most previous studies randomly select a sample size (i.e., the number of single-cell spectra to collect per environmental sample) ^22, 24^, estimate empirically ^25^, or follow lab-specific protocols ^11, 26^. A commonly selected range of SCRS dataset size is 200-1,000 in dependence to the population complexity ^23, 27, 28^ or 20-200 spectra per label in classification-oriented studies ^21, 25, 29^ with a largest reported total sample size of 10759 ^30^. Second, the level of microbial community diversity in different environmental samples varies largely, therefore it is difficult to estimate *a priori*. The optimal SCRS sampling depth for any given system remains unsolved and it limits the standardization of SCRS for its wider applications. This drives the demand for a robust method which statistically validates the sampling depth without knowing the composition and complexity of the microbial community.

Sampling size assessment of SCRS from environmental samples were discussed in previous studies but is often restricted to situational applications. Learning-curve (LC) based technique targeting 5% Bayes error rate was proven effective to investigate proper sample size to train a classifier ^31, 32^; however it is a quite different objective and this method is not suitable for unsupervised applications. Majed et al. (2009) first attempted a practical solution by iteratively sampling and classifying samples, tracking abundance changes of classified categories ^25^. Their relative abundances would be repeatedly calculated, adding a fixed number of spectra each time, until they stabilized above a sample depth threshold. However, its reliance on classification requires a significant amount of both human intervention and domain knowledge to select appropriate discriminating criteria. For example, in their proposed protocol, an exploratory experiment was first carried out to identify 65% biomass (in biovolume) as a functional group of known metabolic traits. The choice of classification criteria which target this majority type of cells therefore was validated. Despite the cost of preliminary experiments, such a dominating microbial group or species is not guaranteed to exist in more complex environmental samples. introduction of an adopted rarefaction-like technique for enabling direct application on continuous feature datasets (e.g., SCRS) without pre-classification, which defined maximum pairwise Euclidean distance as the diversity measure of a sample set, was named as “diversity index (DI)” ^28^. However, two potential issues exist with this DI-based approach. First, the DI of the entire observation dataset is directly treated as reference in sampling size assessment, implicitly regarding it as the population but without further validation. Second, using maximum Euclidean distance to represent the diversity discards all detailed distance distributional structures, potentially causing under-estimation of the true diversity. Thirdly, the definition and parameter for quantifying “diversity” is required, which may not be available such as the case for the SCRS data.

In this study, we propose a new algorithm for sample size assessment that circumvents the disadvantages inherent to previously reported approaches. Our algorithm iteratively increases the number of samples until the optimal sample size is achieved. At each increment, our algorithm measures the distributional difference due to the increase in sample size via *kernel divergence*, a pseudometric that measures the difference between two population distributions. As more samples are collected and observed, the distributional change due to the added samples also converges towards zero. By observing this decreasing trend, our algorithm is capable of predicting the sample size as the change in distribution becomes less than a user-defined threshold. Compared to previously reported approaches, this method has two major advantages. First, no distribution or linear relationship is assumed in the input data; this enables more general and improved handling of datasets when such assumptions do not necessarily hold. Second, it is unsupervised, granting the independence to either classification or other pre-processing which may require extra prior knowledge (e.g., microbial community composition) and human intervention (e.g., selection of classification criteria). This method is validated in a case study using SCRS datasets sampled from eight independent enhanced biological phosphorus removal (EBPR) microbial communities, each obtained from a different North American wastewater treatment facility. To our knowledge, this is the first population-blind sample size prediction and assessment method that has been applied on biospectroscopic datasets. The outcome will facilitate wider, more standardized and more reliable application of advanced analytical technologies such as SCRS for various environmental studies. In addition, since minimal assumption is made with the input data, this approach can potentially be applied on sampling size assessments with any other dataset as long as it only contains continuous features.

## METHODOLOGY

Given a population *Y*, we wish to discover the smallest possible subset of the population that still statistically represent the total population. If we denote the subset as *X* it can be determined by increasing the size of the subset until its internal statistics become stable, i.e., when adding more samples no longer updates the distribution of the subset. Kernel Divergence (*D*_*p*_) allows one to measure the non-linear distribution difference between two distributions. This study leverages this pseudometric to evaluate the statistical change between increments of samples added to the subset. By iteratively tracking this value, the size of the smallest sufficient subset can be determined when the distributional change becomes negligible.

### Kernel Divergence

Inspired by He et al. (2017), we noticed that to reliably measure the distributional differences between two sample sets is critical in sampling size assessment. There are currently many ways of measuring the distributional difference between two populations. Broadly, they can be organized into two categories: F-divergence ^33^ and Integral Probability Metrics (IPMs) ^34^. F-divergence uses the ratio between two distributions to measure their similarity while IPMs use their difference. In general, F-divergence requires the researcher to know the distribution ahead of time. This makes it difficult to compute the divergence given just samples, e.g. the Kullback-Leibler divergence ^35^. Alternatively, IPMs do not require prior knowledge of the sample distributions. Instead, they approximate the distribution directly from the samples. A standard IPM is the Maximum Mean Discrepancy (MMD). It measures the similarity between two distributions by comparing their 1st moment in the Reproducing Kernel Hilbert Space (RKHS). In its original space, it consequently compares all of its moments. Kernel divergence is different in that it uses the 2nd moment in the RKHS, therefore, it is a shift and rotation invariant in RKHS. Instead of comparing the distributions based on their moment, kernel divergence looks at the shape of the data in RKHS as measured by its variance along its principal components. This approach is extremely memory efficient as it allows us to compress the distributional information with a couple of eigenvalues. Any new incoming population can consequently be also reduced to these numbers to compare their distribution difference.

We propose the usage of kernel divergence, motivated by two observations. First, by mapping data onto the RKHS, the non-linear aspects of the data can be captured for analysis^36^. Second, the variances along the principal components (PCs)^37^ of a dataset summarize the shape of the data. **Figure 1** (a) and (b) shows a scatterplot of two populations from the same distributions. Note that PC variances of the distributions in (a) and (b) closely matches each other, indicating a resemblance in the shape of the data. Alternatively,

**Figure 1.**
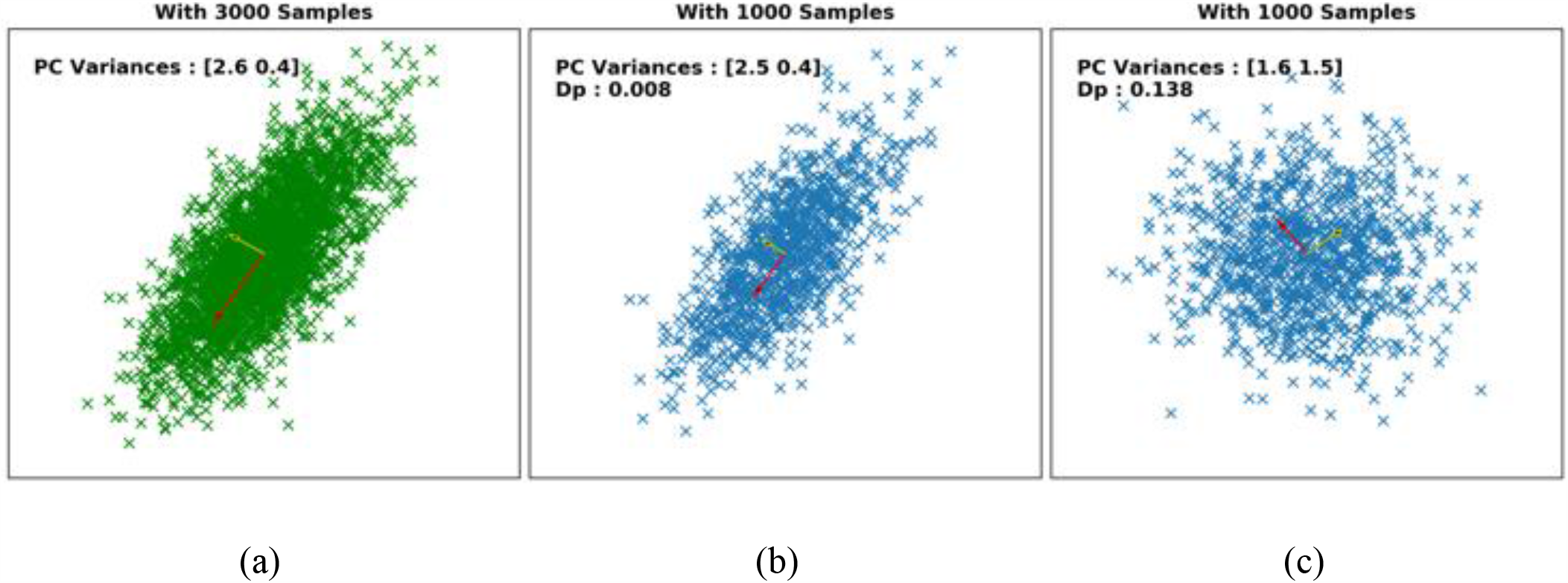
Example of how the principal components can be used to compare distributions. (a) and (b) were generated by an identical population while (a) has three times the number of samples than (b). (c) was generated from a different distribution and exhibited different variation ratios along PC1 and PC2 in comparison to (a) and (b).

**Figure 1 (c)** shows a scatter plot from a population belonging to a different distribution. Note that its variances along the PCs are also noticeably different from that in (a) or (b). Unfortunately, since the PCs can only capture linear relationships, it is no longer an appropriate tool when the data fail to match the linear assumption. Since the data distribution from real applications are commonly unknown, using PCs to compare distributions may not be appropriate for all cases. The idea of Kernel Divergence is to combine RKHS’s ability to capture non-linear relationship with PC’s ability to summarize the data. By first projecting the data into RKHS, non-linear PCs ^38^ can now be used to compare the distributions. Since the concept of PCs is commonly used for the original data space, we will distinguish the PCs in RKHS as the kernel principal components, or KPCs.

There are several advantages in using Kernel Divergence. First, the value computed by the divergence can be treated as a distance between two distributions. It is always a positive value where a *D*_*p*_ = 0 denotes a complete equivalence of the empirical variances along the KPCs. Conversely, a larger *D*_*p*_ also indicates a further distance between two distributions. Second, the value of the divergence *D*_*p*_ has practical meanings as suggested by its theoretical proof. Namely, it denotes the worst-case error along a KPC. For example, if *D*_*p*_ = 0.01, then the biggest difference between two samples along any single KPC is bounded within 1%. Third, and more importantly to practitioners, *D*_*p*_ can be efficiently computed using only a few lines of code with existing open-source software.

### Computing the Kernel Divergence

We define a population of samples as *X* ∈ ℝ^*n*×*d*^ where *n* and *d* denotes the number of samples and the dimension respectively. Let *H* be a centering matrix defined as 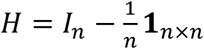 where *I*_*n*_ denotes an identity matrix of size *n* and **1**_*n*×*n*_ ∈ ℝ^*n*×*n*^ denotes a matrix of 1s. Then a centered kernel matrix is defined as *HK*_*X*_*H* where i-th row and j-th column element of the *K*_*X*_ is defined as

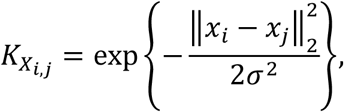

known as radial basis function (RBF) kernel, where exp(·) stands for the exponential function with base *e*, and 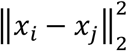 is the squared Euclidean distance between i-th and j-th samples. We chose this kernel function for its flexibility to approximate a wide range of non-linear functions.

Given the definition of a centered kernel, we define *K*_𝔸_ and *K*_𝕊_ as the centered kernels for two population samples of 𝔸 and 𝕊. Let the *m* largest eigenvalues of *K*_𝔸_ be 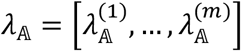, where 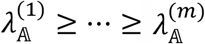, and the *m* largest eigenvalues of *K* be 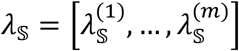, where 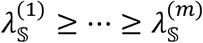. Here, *m* is preferably the number of “major” eigenvalues indicated being before a significant value drop in the eigenvalue spectra plot. Eigenvalues are the variances in the respective eigenvector directions. It is notable that some datasets will exhibit gradually decreasing eigenvalues with no such drop. In such cases, an extra parameter *p*, the percentage of the total variance which we wish to preserve from the population, is introduced to calculate *m*; specifically,

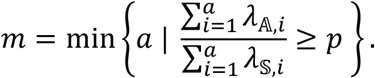

Once the centered kernels of the two populations are calculated, the kernel divergence between *K*_𝔸_ and *K*_𝕊_ is defined as

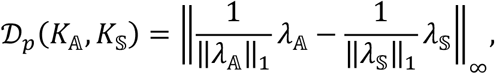

where ‖·‖_1_ and ‖·‖_∞_ are *L*_1_ and *L*_∞_ norms respectively. A brief proof of kernel divergence as a measure is provided in Supporting Information **Proof S1**.

### Sample Size Assessment

Assume that a sample set 𝕊 has already been physically acquired from a population 𝕡. However, knowing only this end-status of sampling process but no *a priori* population composition knowledge for reference, sample size assessment would be almost impossible. To overcome this difficulty, we model the sample acquisition process for better profiling of the sample distribution changes via randomized virtual sampling simulation; while, this modelling step implicitly assumes that the physical sampling experiment (which acquires 𝕊) is unbiased. This will result in a kernel divergence profile at each sampling depth, during acquisition towards the same dataset 𝕊 while in a randomized order. Such simulation can be repeated multiple times for estimation of the mean kernel divergence profile with minimal dependence to the randomization effects. These modelling steps summarize the core algorithm in our assessment protocol, represented as in the pseudo-code below:

1. *A*_1_← randomly select *k*_0_ samples from 𝕊;
2. *A*_*n*+1_← randomly select *k* samples from 𝕊\*A*_*n*_, insert to *A*_*n*_;
3. Calculate kernel divergence *D*_*p*_(*A*_*n*_, *A*_*n*+1_);
4. Repeat (2)-(3) until 𝕊\*A*_*n*_ = ∅;
5. Repeat (1)-(4) multiple times; this results in multiple kernel divergence profiles by acquiring the same dataset 𝕊 in different orders.

For simplicity, at each Step (2), the number of samples drawn is standardized as a fixed number *k*, denoted as *batch size. k* is often determined by the physical analytical system. This is a sensitive parameter to kernel divergence calculation, thus should be determined *a priori* and kept unchanged during a single assessment. For SCRS data, since the Raman system yields single spectrum for each cell at each sampling event, the *k* value is therefore 1. Finally, we determine sample sufficiency of dataset 𝕊 by checking if the “average” kernel divergence profile resulted from the above steps has converged to zero with a pre-defined threshold *t*. Applying the interpretation of kernel divergence, the presence of such point of convergence (POC) identifies a sampling depth at which further addition of *k* more samples (i.e., a *batch*) will no longer significantly update the sample distribution. In other words, the presence of such POC implies that 𝕊 is *sufficient*, otherwise *not sufficient*. The eigenvalue preserving percentage *p* (if used in kernel divergence calculation) and the number of iterations in Step (5) (default: 1000) are the last two adjustable parameters in our proposed protocol, in addition to *k* and *t*.

### SCRS Datasets from EBPR Facilities

SCRS-based phenotypic survey was conducted on 8 different EBPR-related sludge samples, each generating an individual SCRS dataset. Investigating the community composition phenotypic characteristics captured by Raman spectra is conceptually analogous to the operational taxonomic units (OTUs) based survey via 16S-rRNA amplicon sequencing ^23^. Each sludge sample represents the microbial community in the EBPR anaerobic reactor of a different wastewater reuse and reclamation facility (WRRF) across the North America, with various geological, configurations, operational and influent water characteristics (**Table S2**), including 4 conventional EBPR and 4 side-stream EBPR processes. Studies showed that Raman spectra are sensitive to the experimental conditions and instrumental factors therefore all an identical protocol was followed in acquisition of those 8 SCRS datasets to maximize their cross-comparability. Briefly, each sludge sample was independently performed a phosphorus release and uptake kinetics batch test as described by Gu et al. (2008) ^39^. Raman-based phenotypic survey was performed on the sludge extracted throughout the batch test following the preparing and acquisition protocol described by Majed et al. (2008) and Onnis-Hayden et al. (2019) ^40-42^. All spectra were acquired with a 400-1800 cm^-1^ range which is often referred as the “fingerprint range” for various cellular or biomass substances ^6, 23, 43^. All acquired spectra were then preprocessed with LabSpec 6 (HORIBA, 2 Miyanohigashi, Kisshoin, Minamiku Kyoto 601-8510 Japan), for cosmic spike removal, smoothing, background subtraction, baseline correction and vector-normalization. The survey resulted in a total of 922 spectra in eight WRRF-specific datasets labelled from A-H. Two datasets, F (207 spectra from Westside Regional, S2EBPR) and H (214 spectra from Upper Blackstone, conventional EBPR) had larger sample size in comparison to the other six (ranging from 80-89).

## RESULTS AND DISCUSSION

The proposed protocol was first tested with synthetic datasets to demonstrate its sample size assessment performance and results interpretation. Multiple tests with different parameter settings were conducted for discussion on the selection of parameters with respect to the experimental protocol in real applications. Then the model was validated in a case studying with microbial SCRS phenotyping datasets acquired in experiments.

### Test with Synthetic Datasets

The tested synthetic datasets include both simple, linear and non-linear relationships. The dataset construction and the test results are as follows:

#### Dataset construction

The first dataset (Dateset 1) is a series 2-dimensional datasets to test the model’s performance on sample size assessment (**Figure 2** (left)). A total of five sub-datasets were generated from an identical mixture distribution to resemble five independent sampling experiments from a same, infinite-sized population but targeting at different sampling sizes, respectively 20, 40, 100, 200 and 500. The source distribution is composited by four 2-dimensional Gaussian distributions (i.e. *D* = 2, *K* = 4), and is designed to be non-overlapping and linearly separable for better testing our model performance with simple datasets. Therefore, each Gaussian sub-population is centered respectively at (±2.5, ±2.5), with variance of 0.1 on both axes.

**Figure 2.**
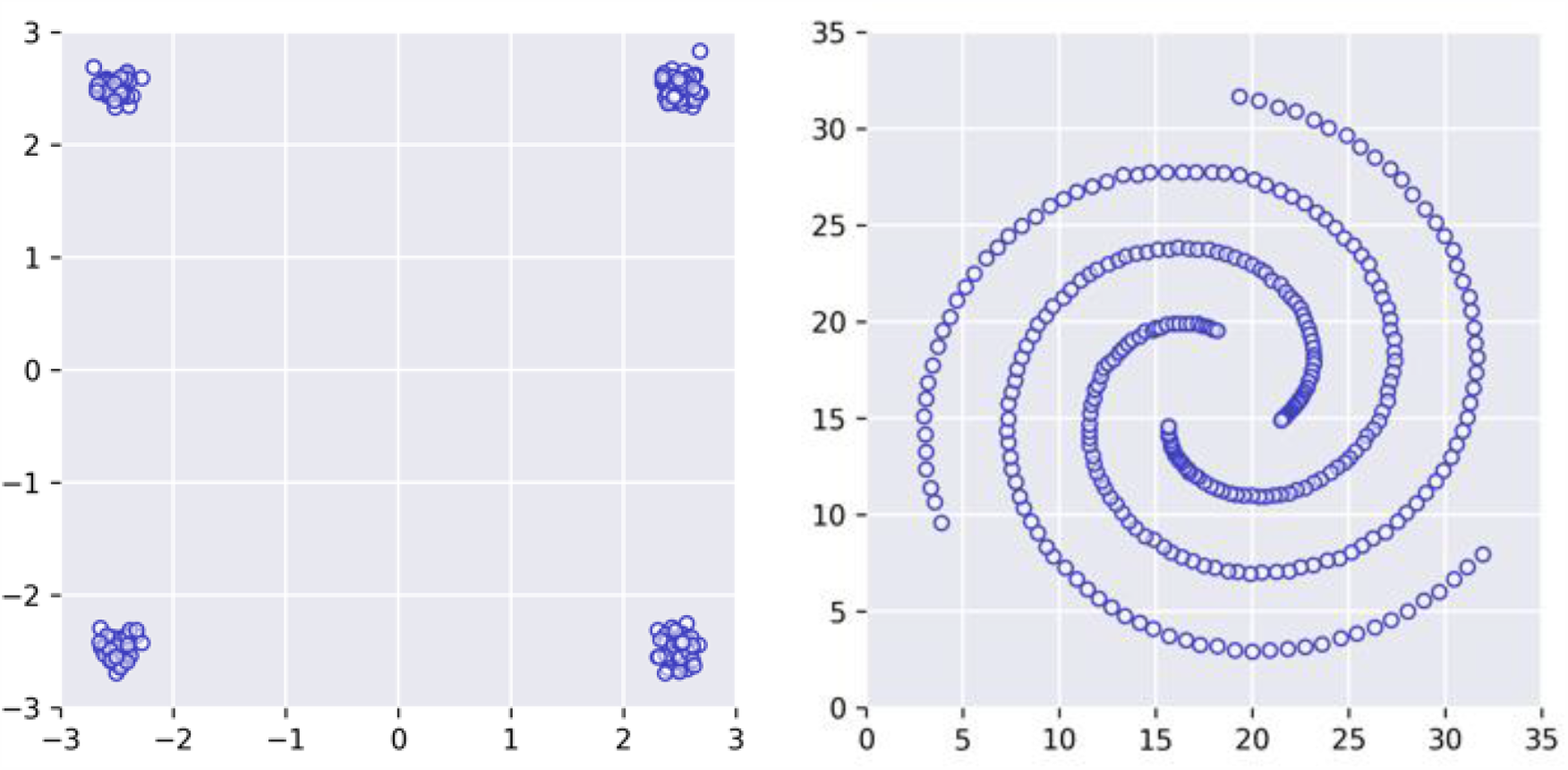
Visualization of synthetic datasets used in performance testing of proposed sample size assessment method using kernel divergence. Dataset 1 (left) is generated with a mixture composed of four 2-dimensional Gaussian sub-populations, each has 25% of the total population and independent variance of 0.1 on both axes. It contains five sub-datasets of various sizes (20, 40, 100, 200 and 500). The 100-sample dataset is shown as a representative. Dataset 2 “spiral” (right) is a 312-sample, 2-dimensional dataset containing three clusters, each forms a non-linear spiral shape ^44^.

Dataset 2 has a spiral morphology ^44^ aimed for testing with complex, non-linear datasets. This 2-dimensional dataset contains 3 clusters in total of 312 samples (*D* = 2, *N* = 312, *K* = 3) (**Figure 2** (right)).

#### Sufficient sampling size assessment

We first demonstrate the calculation and decision making of the proposed sampling size assessment protocol with Dataset 1. The kernel divergence at different sampling depths are shown in **Figure 3**, calculated using empirical parameter settings with batch size *k* = 1 to simulate the one-by-one sampling strategy, and using empirical convergence threshold *t* = 0.01 and 1,000 iterations. We determined *m* = 3 as almost 100% variation fell on the first 3 KPCs (**Figure S2**). The selection of threshold *t* = 0.01 was identified in sensitivity test corresponding to a 0.8% foreign class abundance (will be discussed in detail later). The mean kernel divergence was plotted as solid lines shaded with the standard deviations observed at each sampling depth.

**Figure 3.**
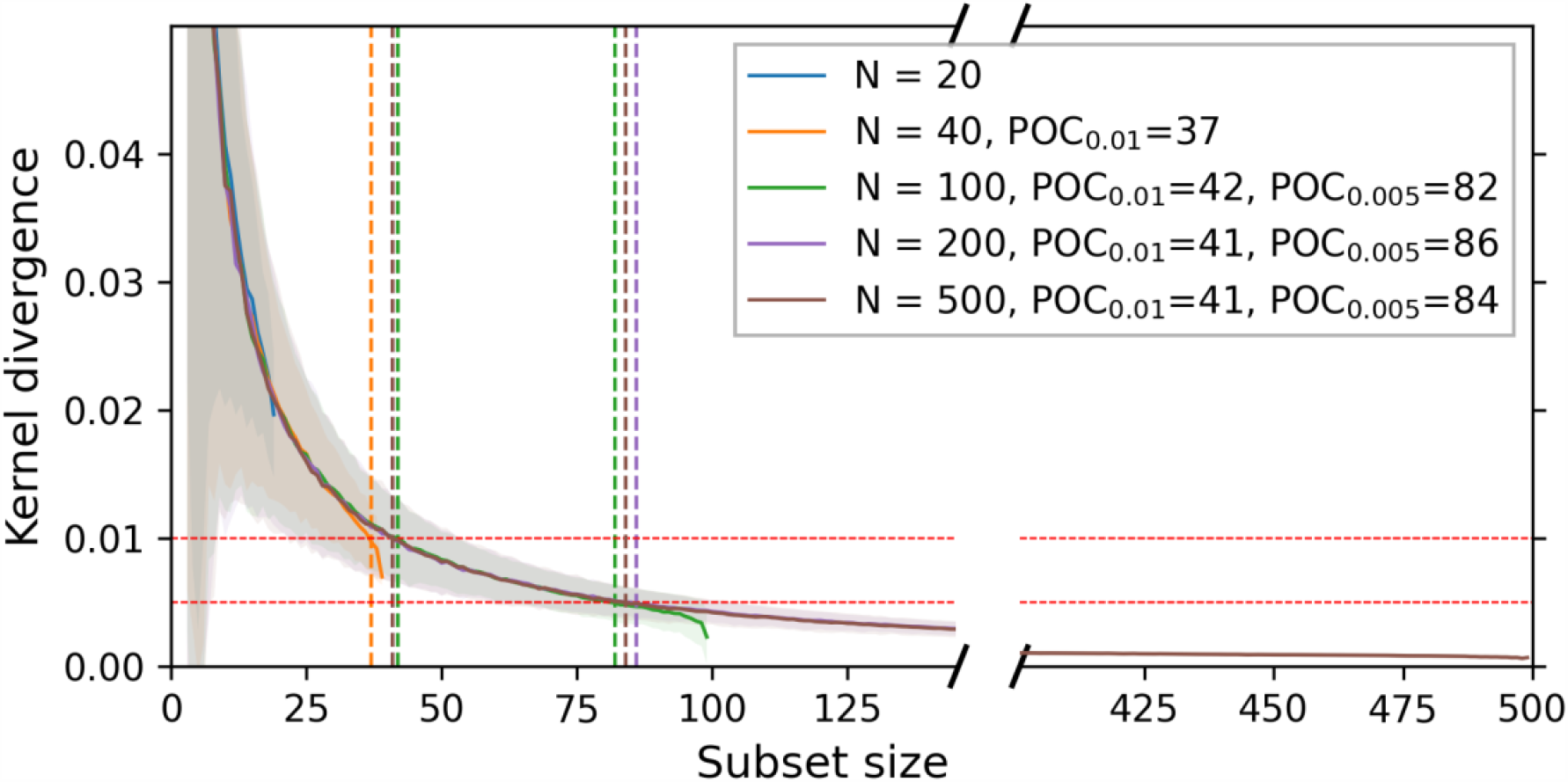
Per sample kernel divergence profiled at various sampling depths simulated independently from each sub-dataset in Dataset 1. All simulations used first 3 eigenvalues ***m*** = **3**, batch size ***k*** = **1**, 1000 iterations and convergence thresholds ***t*** = **0. 01, 0. 005** (shown as horizontal lines). The point of convergence (POC) indicates the minimal sample size required to bound the distributional difference below the indicated threshold, subject to adding one more sample. A lower threshold corresponds to larger sample size.

The closely overlapping kernel divergence profiles indicates that POC estimation is rather robust and consistent, being independent to the dataset size but only the sample distributions. For this data set, the estimated sufficient samples size is around 40 with a targeted convergence threshold of 0.01. As shown in Figure 3, sample size less than 40 (N=20, 40) was not sufficient to meet the targeted POC. Of course, the sufficient sample size depends on the user-chosen POC level. If the target POC threshold is at 0.005, then the sufficient samples size for this data set would be around 85.

#### Assessment with non-linear dataset

A series of similar simulations were conducted on the Dataset 2 “spiral” containing subpopulations that are not linearly separable. All assessment simulations were conducted with 1,000 iterations and three different batch sizes *k* = 1, 2, 5. We chose *m* = 5 eigenvalues which encode approximately 95.5% of total variances along KPCs (**Figure S3**). A convergence threshold *t* = 0.005 was selected for this case, which was identified in sensitivity test corresponding to a 3.3% foreign class abundance. Results are shown in **Figure 4**. The sufficient sample size varied depending on the batch sample size k values. Note that k value is physically determined by the analytical method itself. Here, we demonstrated that the sufficient samples size for targeted POC threshold increased from 59 to 123 as the batch sample size increased from k=1 to k=3.

**Figure 4.**
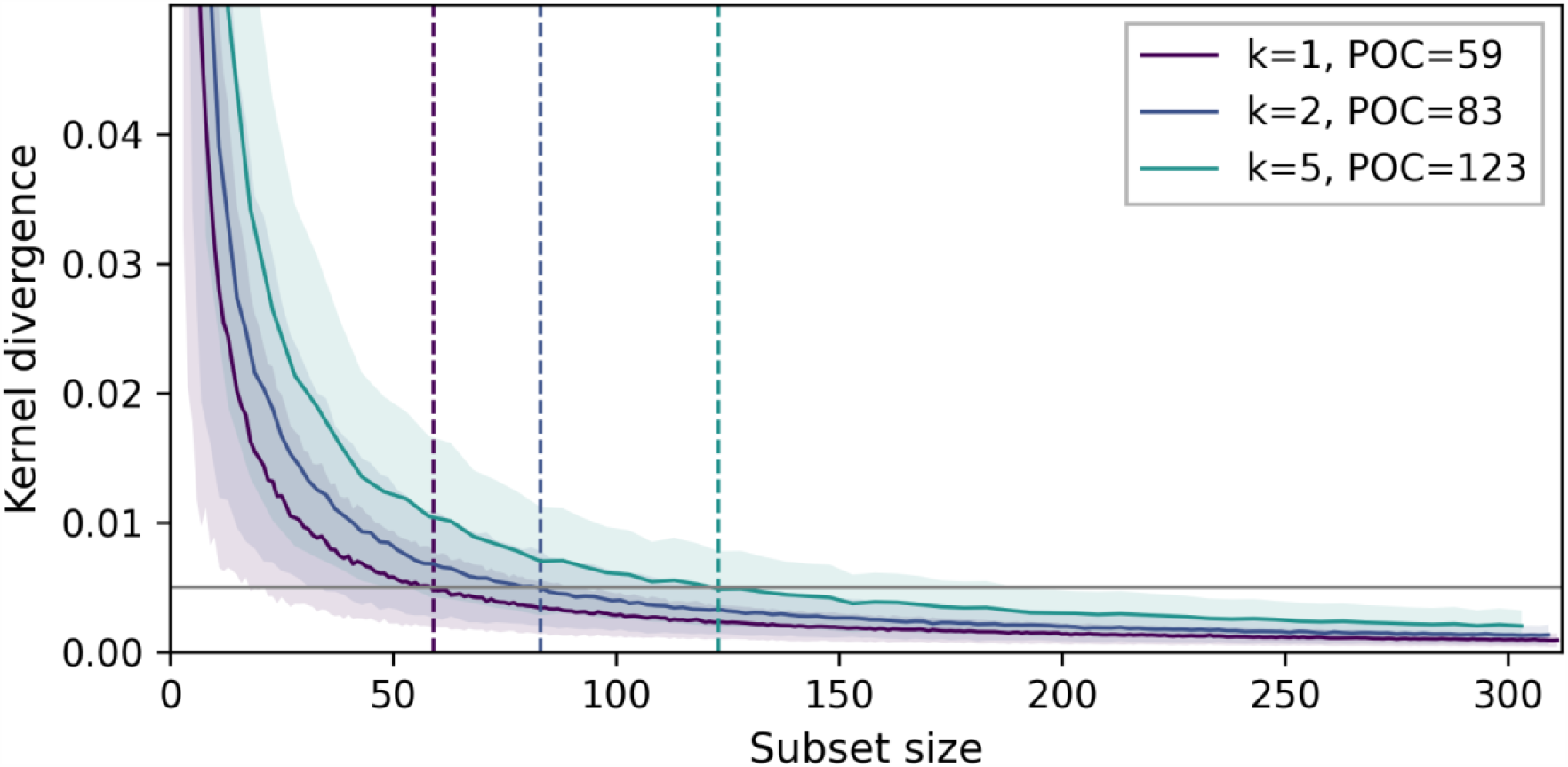
Sample size assessment results with non-linear dataset. Plot shows kernel divergence profiled with different number of samples (1, 2 and 5) per iterative addition (batch size), simulated with Dataset 2 “spiral”. All calculation used first 5 eigenvalues (***m*** = **5**) and were from 1000 simulation iterations. Convergence threshold ***t*** = **0. 005** (shown as horizontal line). The point of convergence (POC) indicates the minimal sample size required to bound the distributional difference below the indicated threshold, subject to each batch size.

**Figure 5** investigates the statistical properties of the POCs in each simulation iterations under different convergence thresholds, which were identified as the last step in each specific simulation iteration having a kernel divergence larger than the given threshold. These results showed that, taking the advantage of RBF kernel, our method can effectively capture the distributional information in a dataset with non-linear aspects. For example, with convergence threshold *t* = 0.005, 2.5%-97.5% quantile of identified POC subsets had estimated class means being within ±2.03 of the actual population mean on both dimensions. And, the estimated class abundances were within ±7.0% of actual abundance. At 25%-75% quartile, error of class means, and abundances were ±0.72 and ±2.7% respectively. A smaller threshold would result in more parametrically accurate POC subsets; however, it requires more samples to achieve reliable POCs. The test showed that the POC subset sizes had 25%-75% quartile respectively from 47-61 with *t* = 0.01, 102-124 with *t* = 0.005 or 307-312 with *t* = 0.001. In addition, incorporating the radial basis function (RBF) kernel enabled the kernel divergence to be appropriate in measuring dissimilarity between two datasets with non-linear aspects, and therefore our method is generally applicable in other datasets.

**Figure 5.**
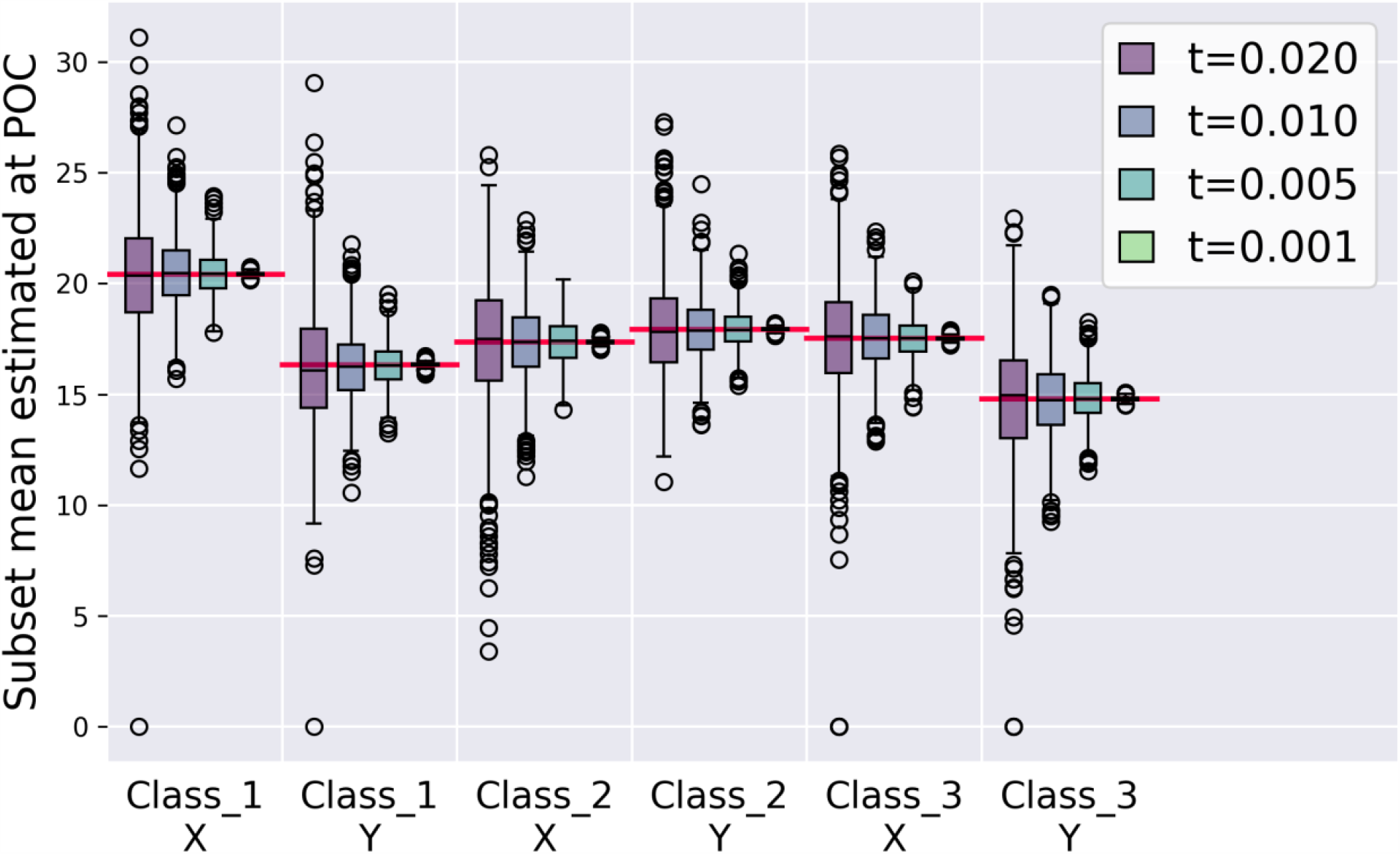
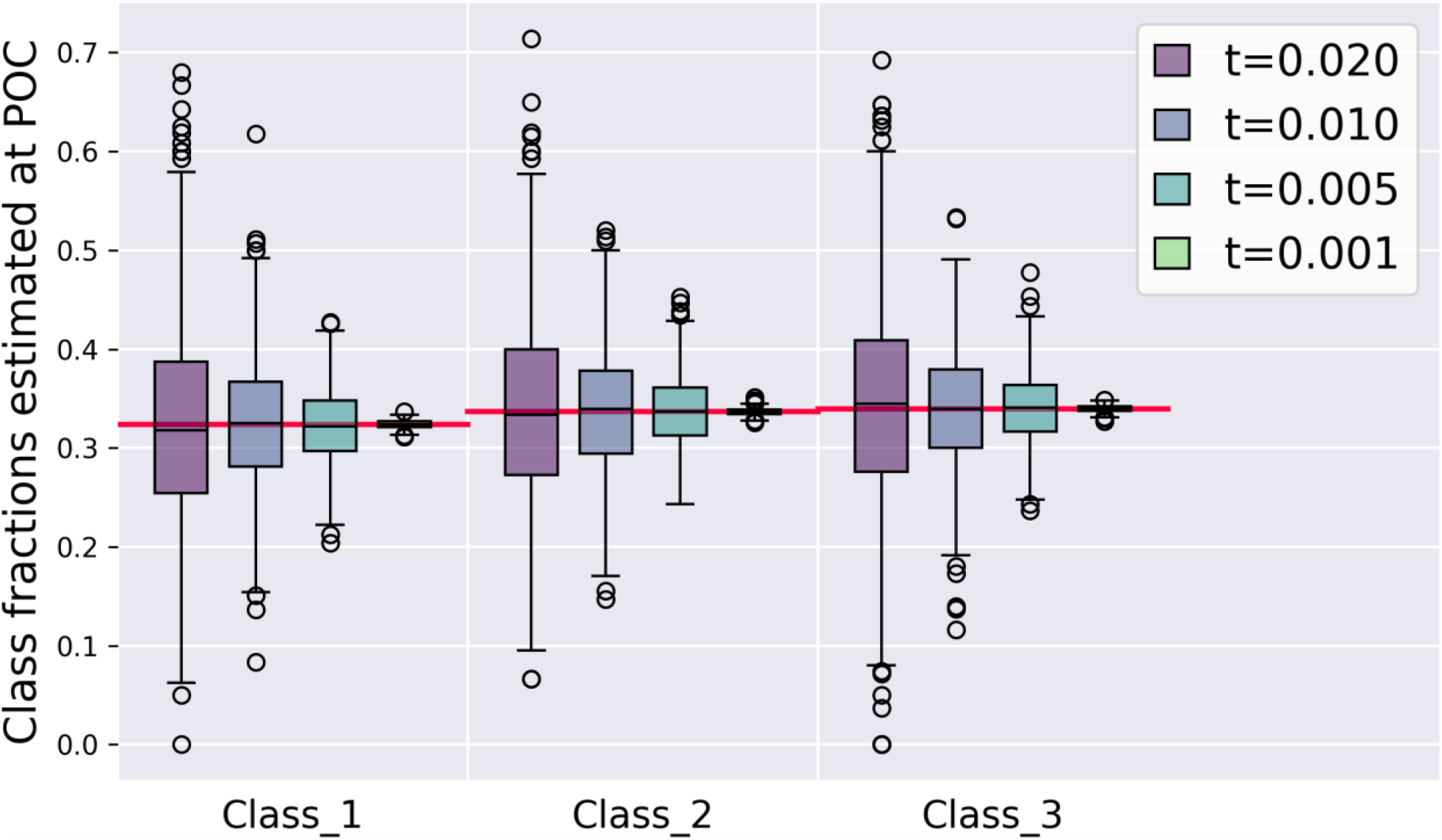
Box plots of mean (top) and fractions (bottom) for each class in the subset selection at each individual POC with different convergence threshold ***t***. from the 1,000 simulation iterations. The simulation POC is determined as the last step in specific sampling simulation that had a kernel divergence larger than the given threshold ***t***. The horizontal red lines indicate the ground truth calculated from the whole “spiral” dataset.

#### Selection of convergence threshold t

Results in **Figure 5** showed the relationship and potential impact to the mean and abundance estimations of each class associated with different convergence threshold criteria. As the threshold pre-determines the resolution of two datasets being asserted “different”, it also relates with the minimal abundance of identifiable classes. To reveal this effect, we chose one class from the original datasets, Dataset 1 and 2, (referred to as “the foreign class”), and randomly delete a sub-selection of its samples. This creates artificial datasets with the foreign class at varying abundances. With random repeats and the dataset with no foreign class as a reference, we could estimate the minimal abundance of the foreign class to exhibit a significant change in kernel divergence under chosen threshold. Note the choice of foreign class has minimal impact to the results due to the high symmetry. The results in **Figure 6** showed that under a same kernel divergence threshold level, the size of detectable foreign class varies largely depending on the parent dataset. The simpler dataset (Dataset 1) requires less samples to achieve the same kernel divergence level comparing to the non-linear Dataset 2. For example, mixing foreign class at abundances of 0.4% and 0.8% leads to 0.005 and 0.01 kernel divergence; while 3% and 9.5% abundances of the foreign class would be required respectively with Dataset 2. Considering both the effect of precision, resolution and dataset complexity, we suggest that *t* ≤ 0.01 would be adequate empirically in general, and *t* ≤ 0.001 is considerable when a high accuracy is desired.

**Figure 6.**
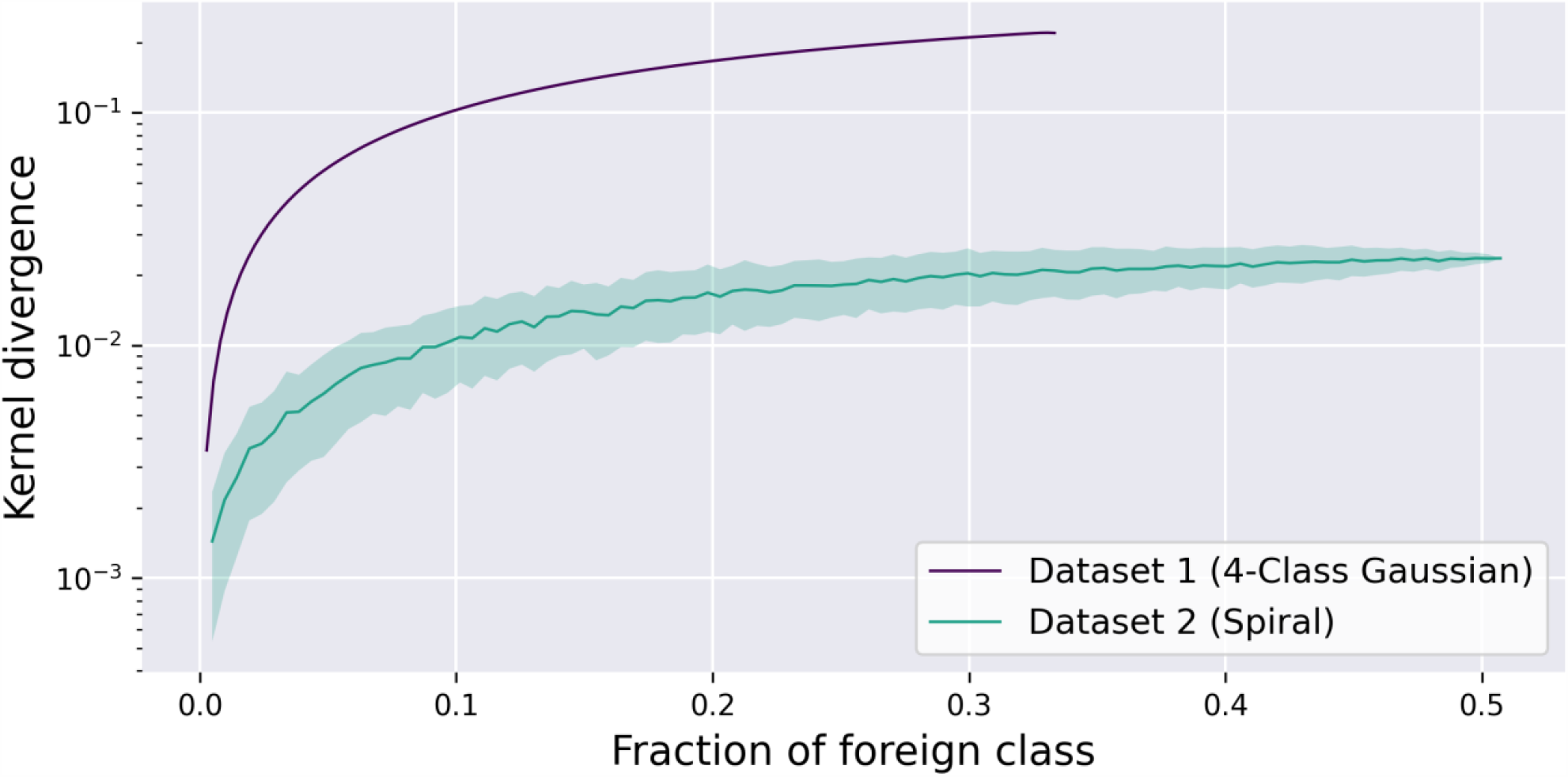
Kernel divergence sensitivity test with varying the abundance of one of the classes (referred to as “foreign class”). All kernel divergences were calculated in reference to the absence of the entire foreign class. The lines and shades show the mean and standard deviation respectively estimated by random sub-selection of samples in the foreign class. The standard deviation in the Dataset 1 profile is minimal and barely visible.

### Case Study: Sample Size Assessment for SCRS Phenotyping Datasets

After performance evaluation on two synthetic datasets with or without linear relationship, the proposed algorithm and method were applied to investigate sampling size requirements with single-cell Raman spectroscopic dataset retrieved from eight microbial communities representing eight different wastewater reuse and reclamation facilities (WRRFs) located in North America ^27, 42^. Details on these EBPR facilities, sampling and SCRS data acquisition were described in the methods section and in supporting information.

#### Sampling size assessment

Kernel divergence profiling simulations were conducted independently on individual SCRS-based phenotyping dataset, using parameter *k* = 1, *t* = 0.01, 0.005 and 1000 permutations. Sample batch size k=1 since our data was obtained using single-cell Raman microspectroscopy at single cell resolution ^27^. As no clear group of major eigenvalues can be identified, we used parameters *p* = 96% in calculations (**Figure S4**).

As shown in **Figure 7**, the sufficient sample size based on each individual kernel divergence profile for the eight EBPR communities ranged from 26-39 under a resolution threshold of *t* = 0.01. The parallel assessment under a higher resolution by lowering the threshold of convergence to *t* = 0.005 revealed an increased sample size range from 47-71 among the EBPR plants. These were below the empirical reliability checking criteria discussed previously (POC: *N* < 0.85). This range are consistent with previously reported values (60-65 samples) proposed by Majed et al. (2009) via an alternative rationale 25.

**Figure 7.**
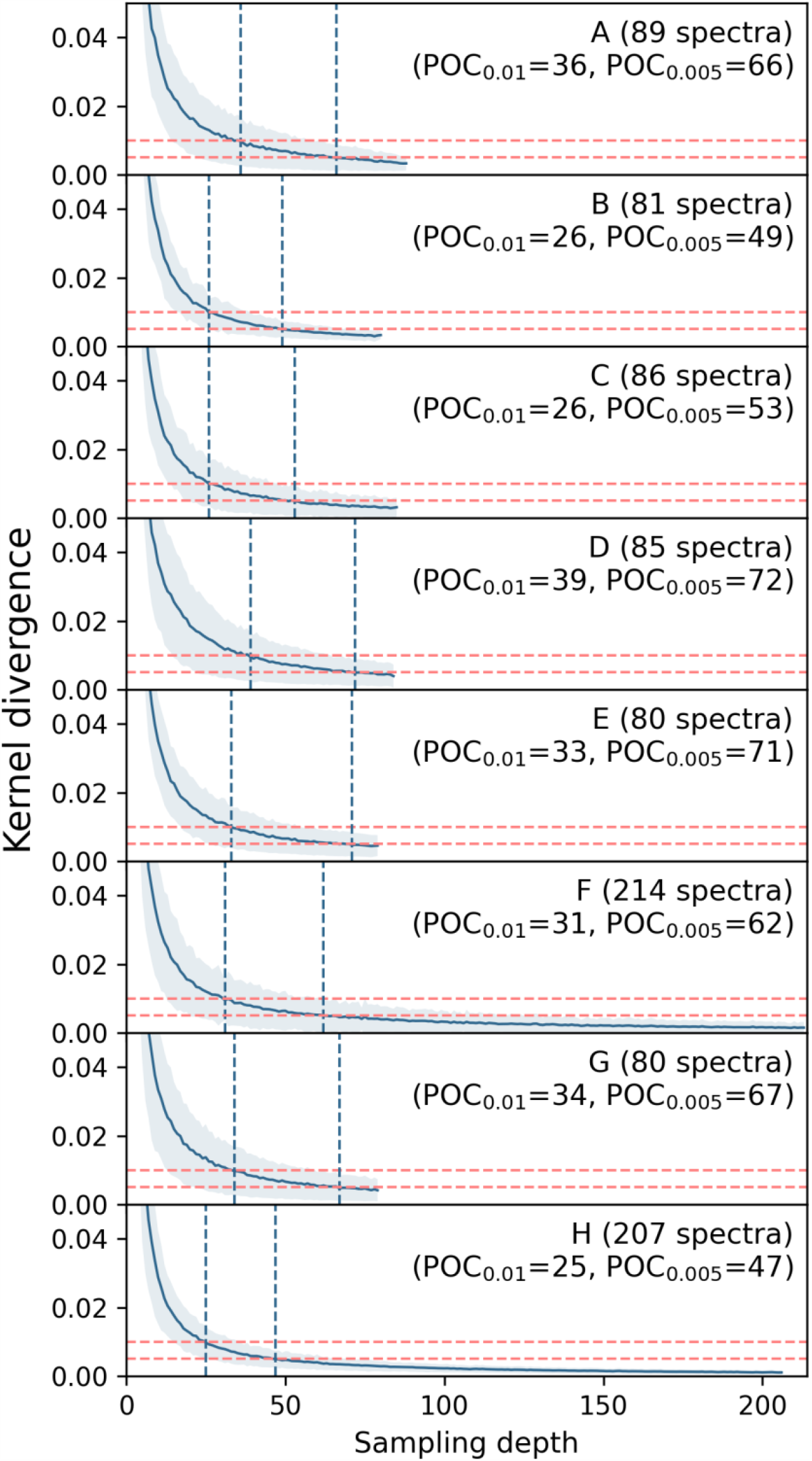
Kernel divergence profiles versus sampling depth, simulated independently on 8 single-cell Raman spectroscopic (SCRS) microbial phenotyping datasets, from 8 individual full-scale enhanced biological phosphorus removal (EBPR) systems in different wastewater reuse and reclamation facilities (WRRFs) located across North America. Simulation parameters were eigenvalue preserving percentage ***p*** = **96**%, 1000 iterations, convergence threshold ***t*** = **0. 01, 0. 005** (shown as the two horizontal lines) and batch size ***k*** = **1** for single-cell Raman microspectroscopic method. The point of convergence (POC) indicates the minimal sample size required based on the targeted convergence threshold ***t*** and batch size ***k***.

#### Investigation of the convergence threshold

The sensitivity test shown in **Figure 8** further investigated the convergence thresholds towards a more physical and practical interpretation, by evaluating the maximum abundance level of sample clusters that could be potentially missed with varying thresholds. Operational phenotypic units (OPU) clustering was first carried out individually to identify cluster groups in each dataset, using correlation distance, average linkage as described by Li et al. (2018) ^23^. **Figure 8** then shows the maximum kernel divergence observed when randomly removing one identified OPU cluster which is below a targeted abundance level. The results indicate that for 7 out of the 8 datasets, using a threshold of 0.01 captures all OPU clusters of more than 5% abundance; and using a threshold of 0.005, this resolution can be improved to 2%. Therefore, we conclude that for these 8 datasets from EBPR communities, the sample sizes were sufficient to capture all OPU clusters of at least 5% relative abundance, and among them, 5 date sets were further sufficient at capturing OPUs with 2% relative abundance.

**Figure 8.**
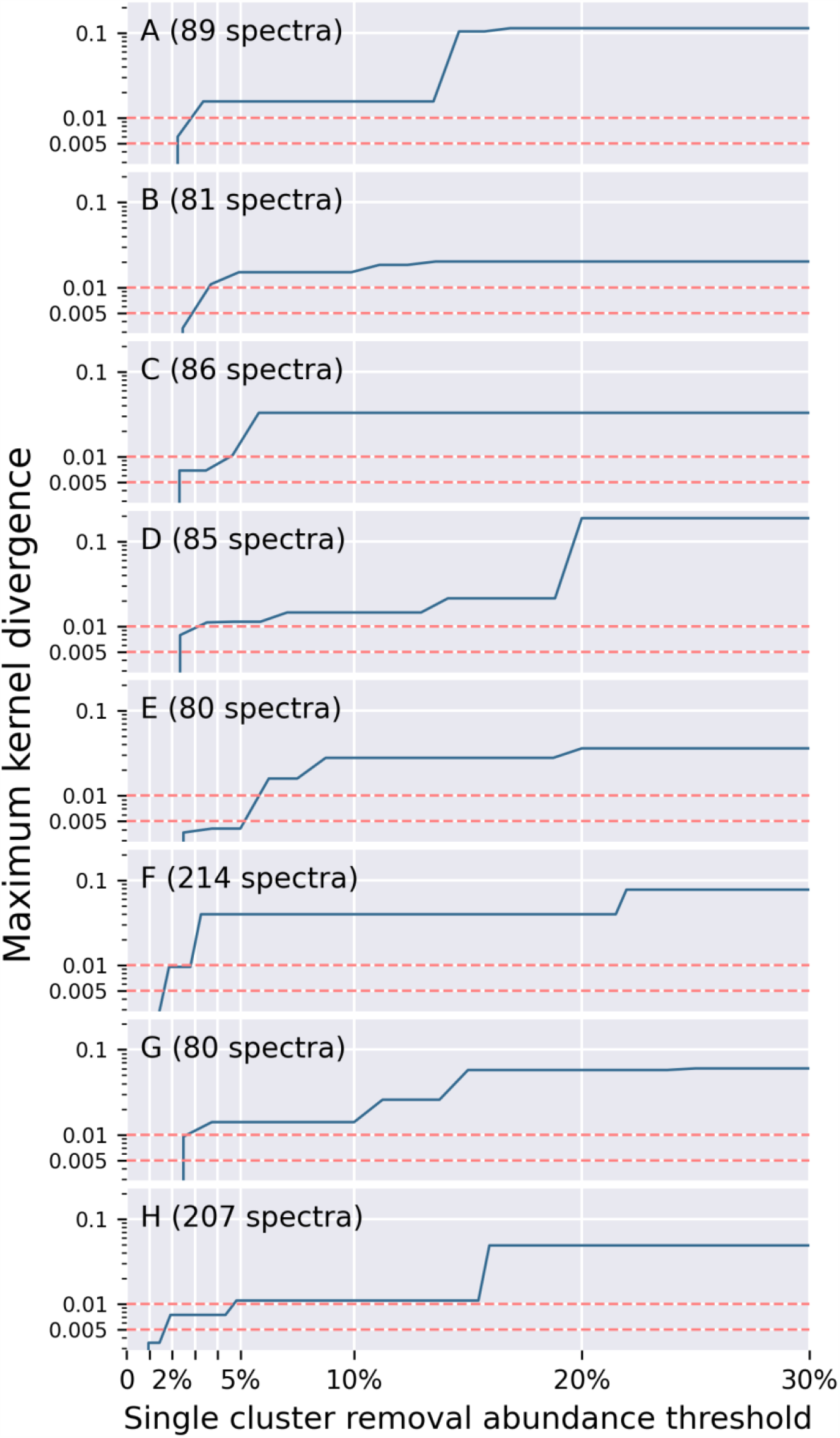
Sensitivity test of kernel divergence convergence over single OPU cluster abundance. The kernel divergence values were calculated by randomly removing a single OPU cluster not exceeding an abundance threshold (x-axis). The maximum kernel divergence calculated at each abundance threshold was shown on the y-axis. The OPU clusters were identified using correlation distance and average linkage as described by Li et al. (2018) ^23^ using a same cut-off at 0.7. The results indicated the relationship between OPU resolution and choice of convergence threshold ***t***. For example, a point x=5%, y=0.01 means that using a threshold ***t*** = **0. 01** allows at most one OPU up to 5% abundance being ignored. Such assessment results may be specific to each dataset.

It is noticed that the identified sample sizes for the same target POC threshold among the eight EBPR communities were rather comparable, indicating an intrinsic “similarity” of the microbial phenotypic “richness” in these full-scale EBPR systems in North America. The correlation between SCRS-based phenotypic clusters (i.e. OPUs) and their phylogenetic OTUs, with underlying implications of the discriminative power of Raman spectrum features for discerning cells at various taxonomic levels (i.e. species, strains etc.), is still under investigation. Promising cell identification at strain level have been reported (ref.). The comparable kernel divergence profiles and narrow range of minimal sample size for a given targeted POC threshold for the 8 EBPR microbial communities suggested that these engineered wastewater treatment systems may have similar microbial phenotypic diversity measurements. How the phenotyping profiles correspond to their phylogenetic composition are yet to be revealed and is beyond the focus and scope of this study.

We also compared our results with another prior-knowledge independent method proposed previously by He et al. (2017) ^28^. The diversity measure of a given sample set, named as “diversity index (DI)” resides as the core concepts in He’s assessment protocol, which was defined as the maximum pairwise Euclidean distance within the sample set. Through repeated virtual sampling experiments in a similar process, the “average DI” at each sampling depth was estimated, then plotted as shown in supplementary **Figure S6**.

Finally, two sample size guidelines can be determined according to the decision criteria proposed by He et al. (2017) ^28^:

1. 9-15 samples as “minimal size” identified when DI change per increasing sample depth by one is less than 0.01 of maximal (i.e. dataset-wise) DI;
2. 9-25 samples as “safe size” identified when average subset DI reaches 90% of maximal (dataset-wise) DI.

These results were significantly smaller than the values identified by our protocol, indicating DI led to a less diverse estimation in comparison to kernel divergence. Two potential reasons may have contributed to different performances of the DI-based and kernel divergence-based protocols. First, DI ignores details of the sample distance distribution but only its maximum, while kernel divergence utilizes comprehensive information of the entire distance matrix. Therefore, DI-based calculations will probably make false conclusions when two sample sets have different sample distributions but rather similar DI values. Second, a sole reliance on the maximum distance also increases its sensitivity to the presence of outliers; therefore, appropriate and sophisticated outlier removal techniques might also be necessary in real applications. Comparing to kernel divergence-based protocol, the DI-based method was likely underestimating the true diversity and complexity in our SCRS phenotyping datasets, resulting in reduced reliability.

One of the main challenges in wider application of new emerging high-resolution technologies for profiling and characterization of complex environmental systems, such as SCRS and cell imaging, is the standardization of the experimental protocols and data analysis such as the optimal sampling size with the consideration of both time and resources cost and information sufficiency. There is no widely accepted approach and method for determining the sufficient sampling size on environmental datasets without pre-classification. We proposed and validated a sample size assessment protocol using kernel divergence, a novel dissimilarity measure at the dataset-level, which is a more comprehensive and systematic quantitative comparison between two observation datasets. More importantly, our proposed method enables the decision on sampling size without prior knowledge of the diversity and complexity of the system. This property is especially powerful as demanded by *de novo* studies with environmental samples. In addition, our proposed method has no restrictions on the input data as long as it contains continuous features. In particular, it can capture data with linear and non-linear relationships. All these generalities profit expansion to further potential applications. First, it provides a universal standard to compare the sampling size determining criteria among different experiments in different labs, contributing to more reliable, comparable and reproducible studies using similar single-cell technologies. In addition, robust cross-comparison among different experimental protocols could be validated as well. Second, we believe that the proposed sampling size assessment approach can be easily generalized to dataset generated from other analytical technologies. Potential examples include Raman-based spectral histopathological assessments, validating gating strategies in flow cytometry and collecting comprehensive cellular imaging library based on visual or morphological measurements.

## Acknowledgement

This work was supported by the United States USDA-NIFA (Grant 2019-67013-29364), NSF IIS (IIS-1546428) and Water Environment Research Foundation (Grant No.). The authors acknowledge to the WERF S2EBPR (U1R13)research team and all wastewater reuse and reclamation facilities participated and provided samples to this project.

## ASSOCIATED CONTENT

### Supporting Information

**Proof S1**. Proof for kernel divergence properties.

**Figure S2**. Eigenvalue spectra plot for the 4-cluster Gaussian dataset (synthetic Dateset 1).

**Figure S3**. Eigenvalue spectra plot for the “spiral” dataset (synthetic Dateset 2d).

**Figure S4**. Eigenvalue spectra plot for the Upper Blackstone and Westside Regional datasets.

**Table S5**. Supplementary information of WRRFs from which the SCRS datasets were sampled.

**Figure S6**. Comparative analysis to the minimal sampling depth and safe sampling depth simulated following the approach proposed by He et al. (2017).

## AUTHOR INFORMATION

### Author Contributions

‡These authors contributed equally.

