## supporting information for "Machine Leaning-based Determination of Sampling Depth for Complex Environmental Systems: Case Study with Single-Cell Raman Spectroscopy Data in EBPR Systems"

**Figure S2.** Eigenvalue spectra plot for the 4-cluster Gaussian dataset (synthetic Dateset 1).

**Figure S3.** Eigenvalue spectra plot for the “spiral” dataset (synthetic Dateset 2d).

**Figure S4.** Eigenvalue spectra plot for the Upper Blackstone and Westside Regional datasets.

**Table S5.** Supplementary information of WRRFs from which the SCRS datasets were sampled.

**Figure S6.** Comparative analysis to the minimal sampling depth and safe sampling depth simulated following the approach pro-posed by He et al. (2017).

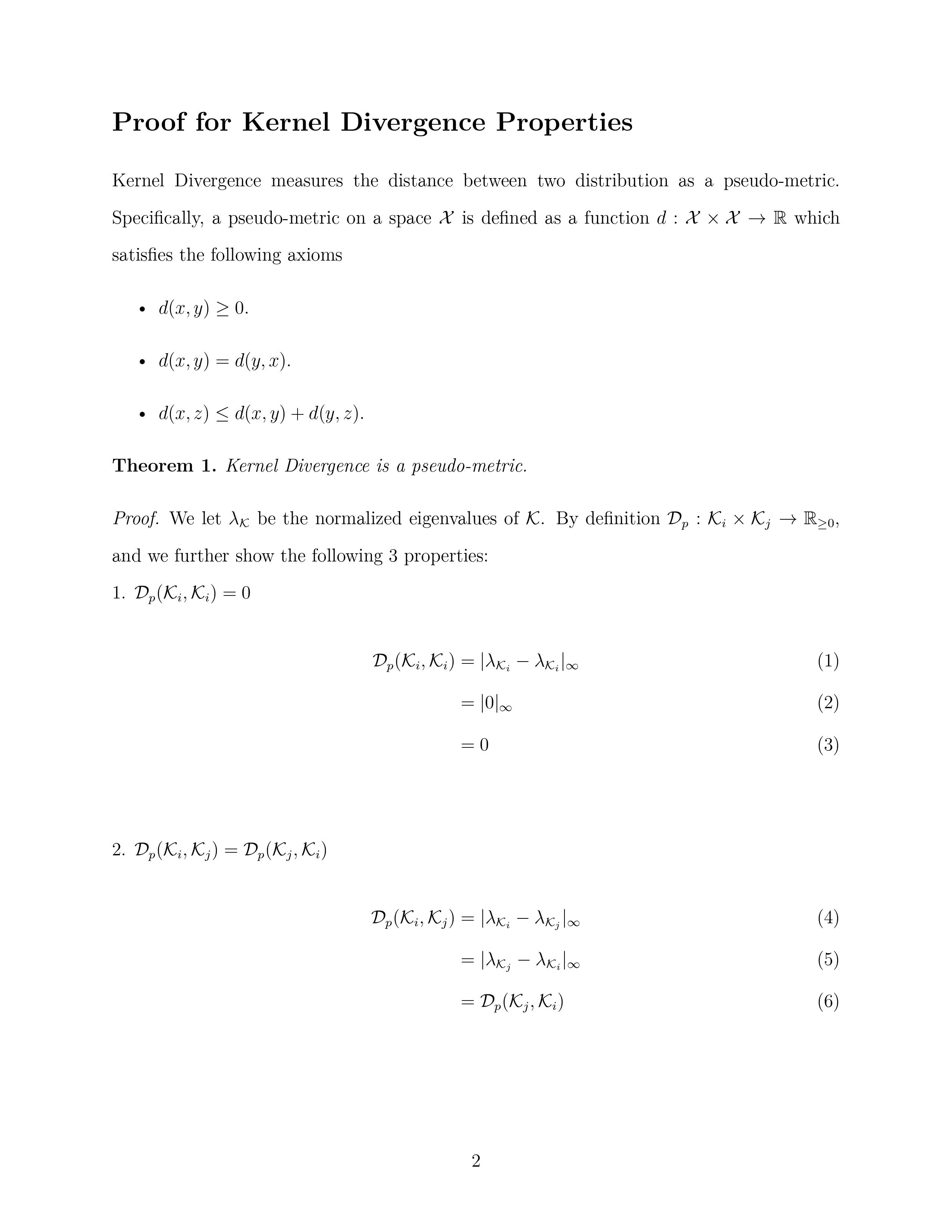

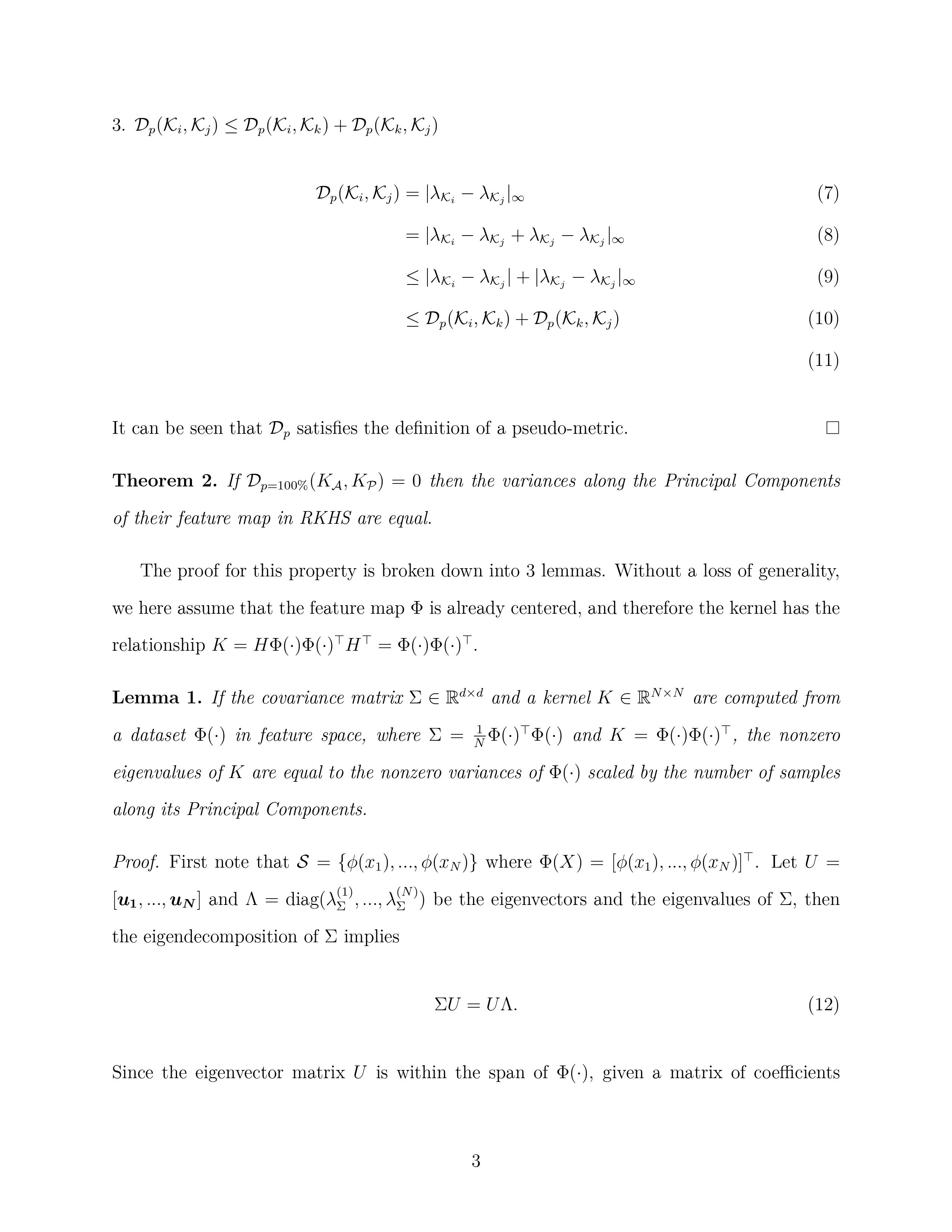

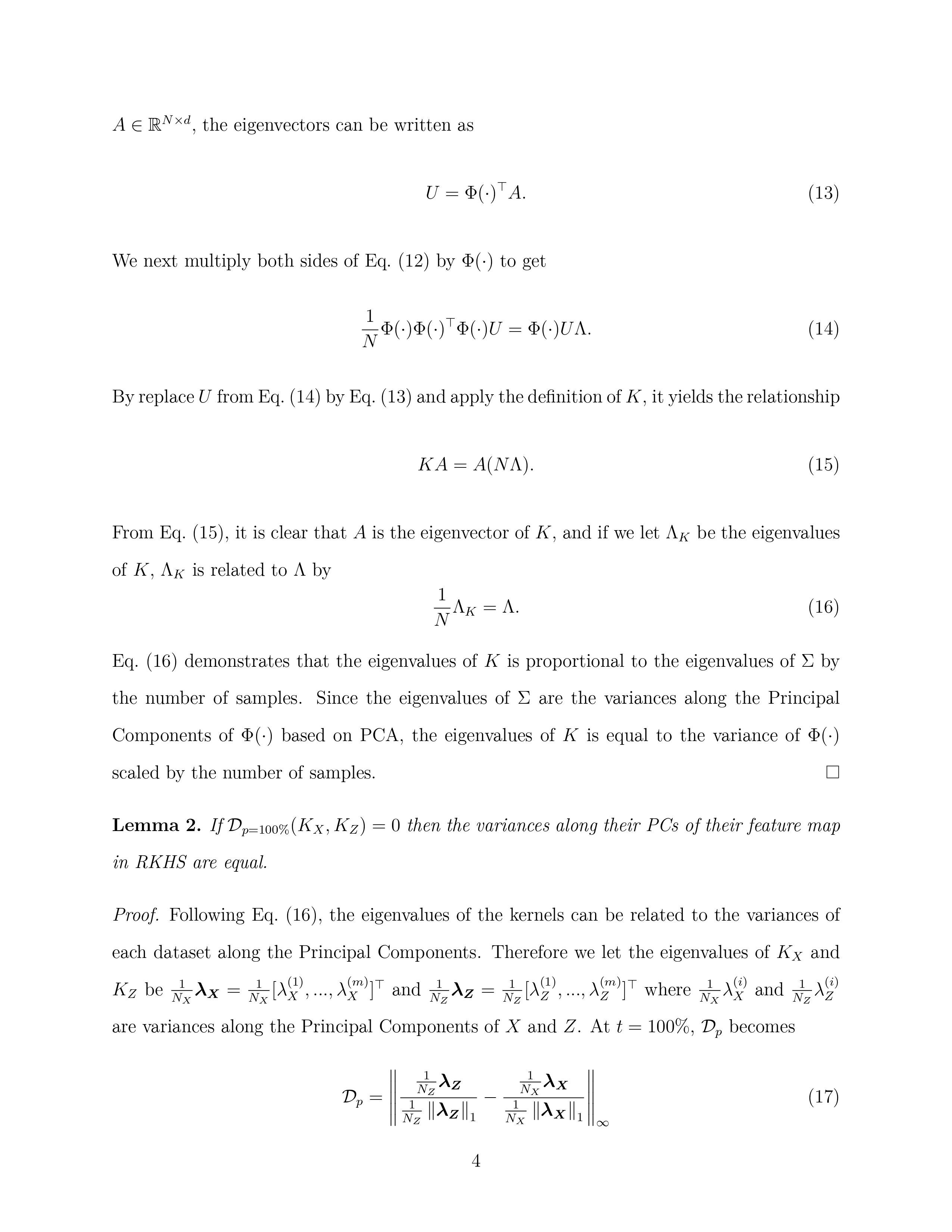

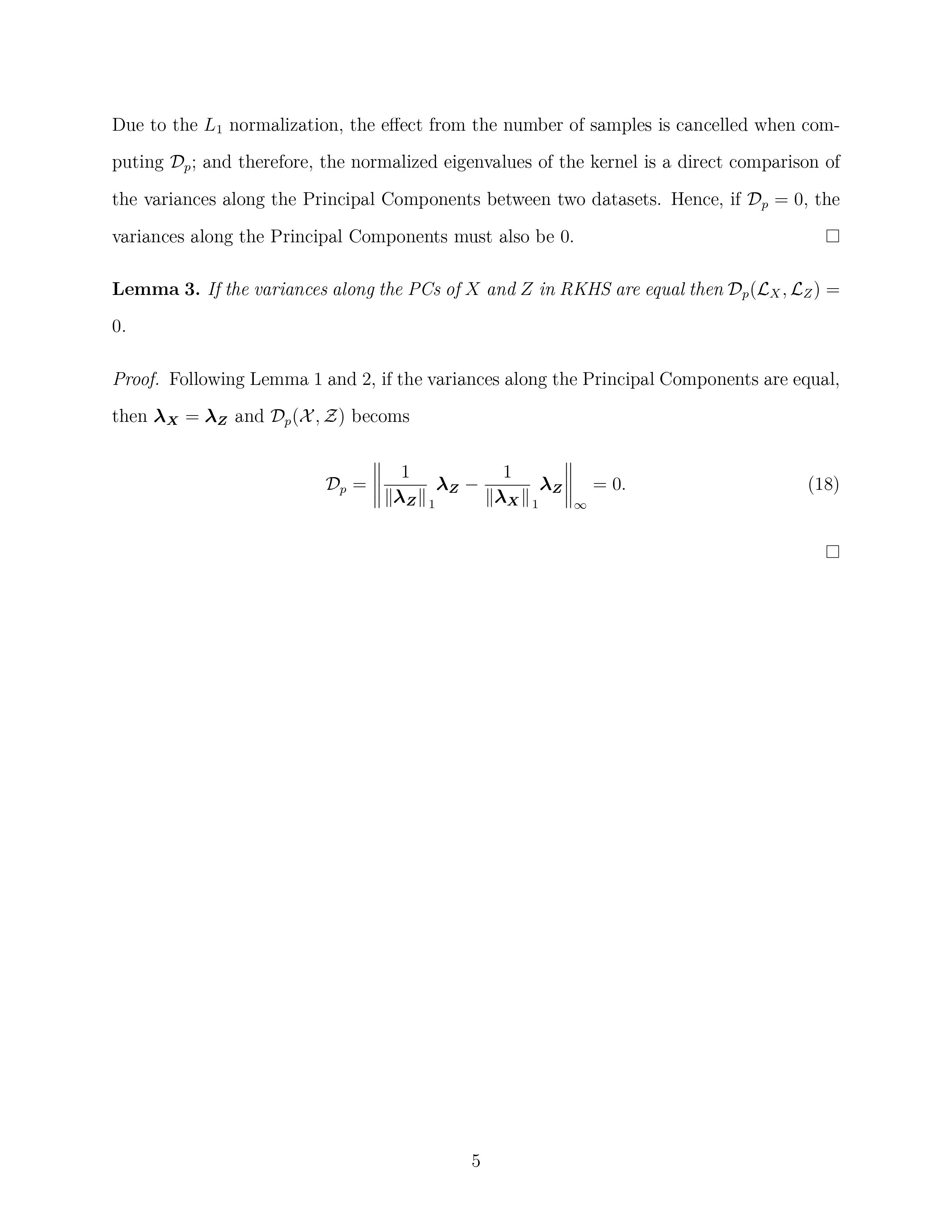

Below **Figure S2** was used to determine the number eigenvalues preserved for kernel divergence calculations, with the 4-cluster Gaussian dataset (synthetic Dataset 1). A big value drop was identified at the 4th eigenvalue. All eigenvalues after this drop were negligible comparing to the first three.

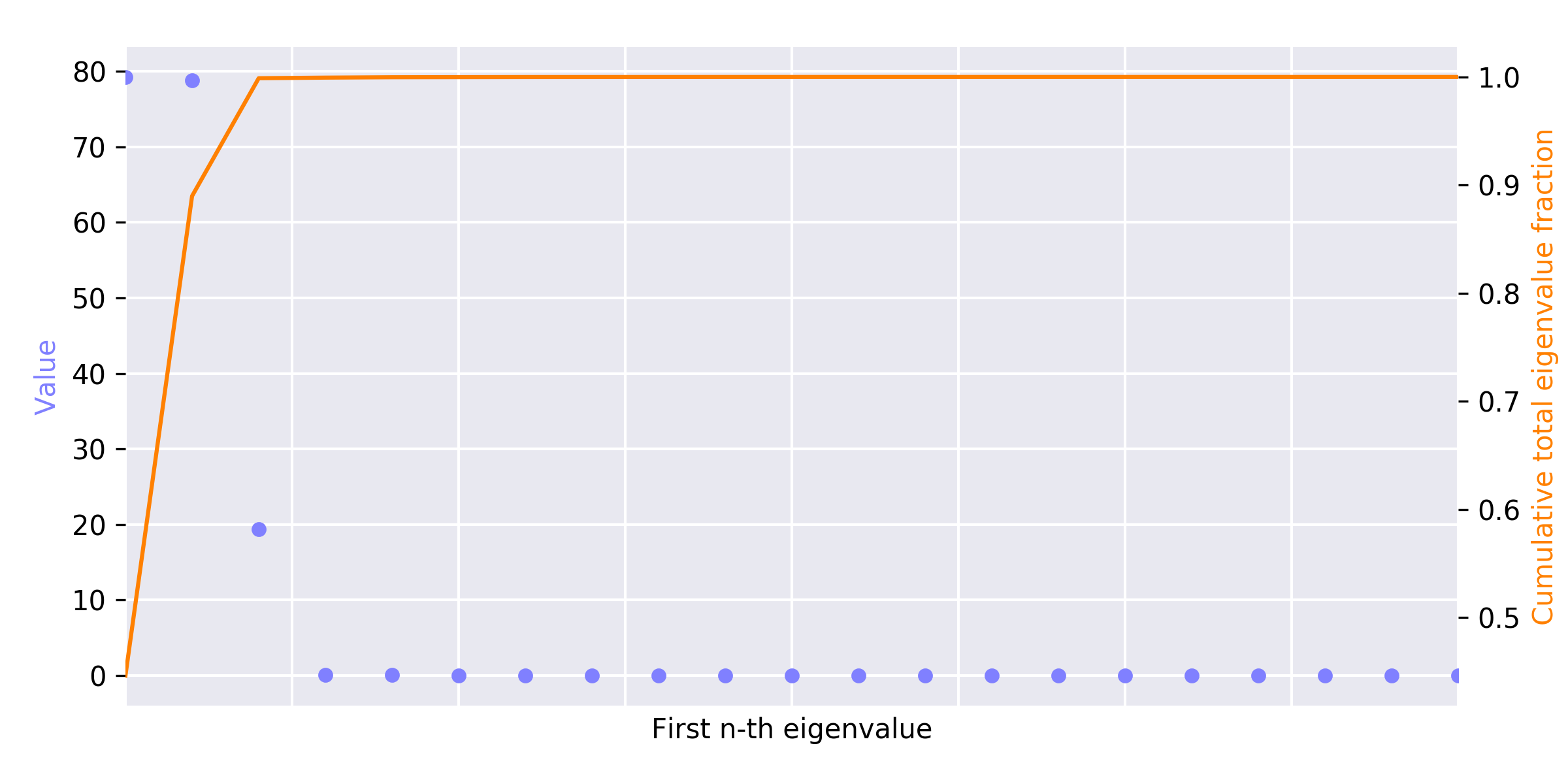

**Figure S2.** Eigenvalue spectra plot for the 4-cluster Gaussian dataset (synthetic Dateset 1). The eigenvalues are sorted in descending order and only the first 20 are shown from the $N=500$ dataset as its representation. The first 3 eigenvalues encode almost all variances on the kernel principal components (KPCs) there fore we identified the $m=3$ in kernel divergence calculations.

Below **Figure S3** was used to determine the number eigenvalues preserved for kernel divergence calculations, for the “spiral” dataset (synthetic Dataset 2). A big value drop was identified at the 6th eigenvalue, and the first 5 eigenvalues encoded 95.5% of the total variances.
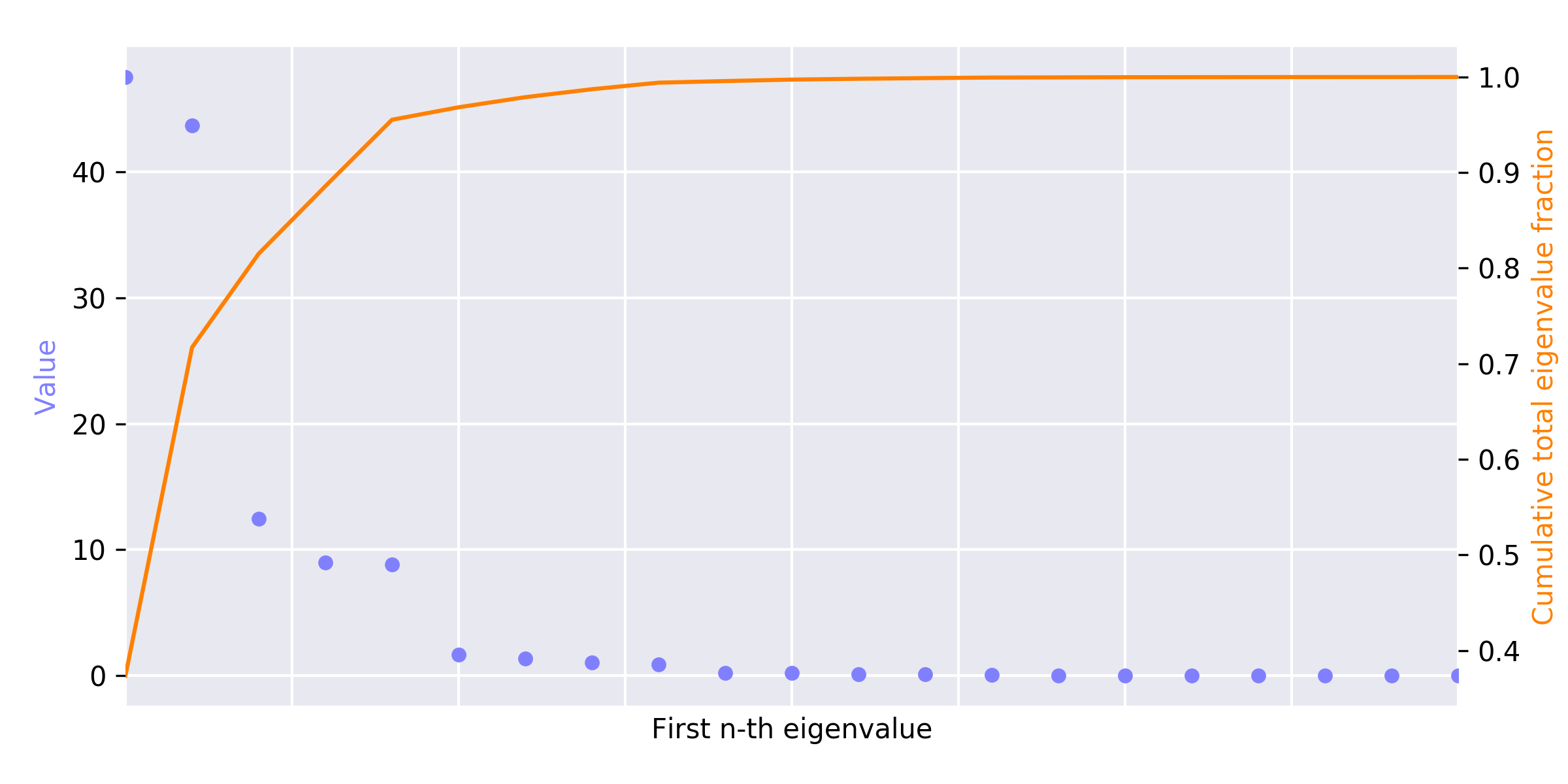

**Figure S3.** Eigenvalue spectra plot for the “spiral” dataset (synthetic Dateset 2). The eigenvalues are sorted in descending order and only the first 20 are shown. The first 5 eigenvalues majority (95.5%) of the variances on the kernel principal components (KPCs) therefore we identified the $m=5$ in kernel divergence calculations.

Below two figures (**Figure S4**) were used to determine the number of eigenvalues used in kernel divergence calculations, for the 8-WRRF SCRS datasets in case study section, showing the two largest SCRS datasets, namely Upper Blackstone (Label F) and Westside Reginal (Label H) as representatives. Unlike synthetic datasets, these more complicated and diverse datasets exhibited no “big drop” in their eigenvalue spectra. Therefore, we use 96% as a cut-off threshold, i.e. we preserved first certain number of eigenvalues that encoded 96% of total variance and ignored all the rest. Note that this number may vary between datasets.

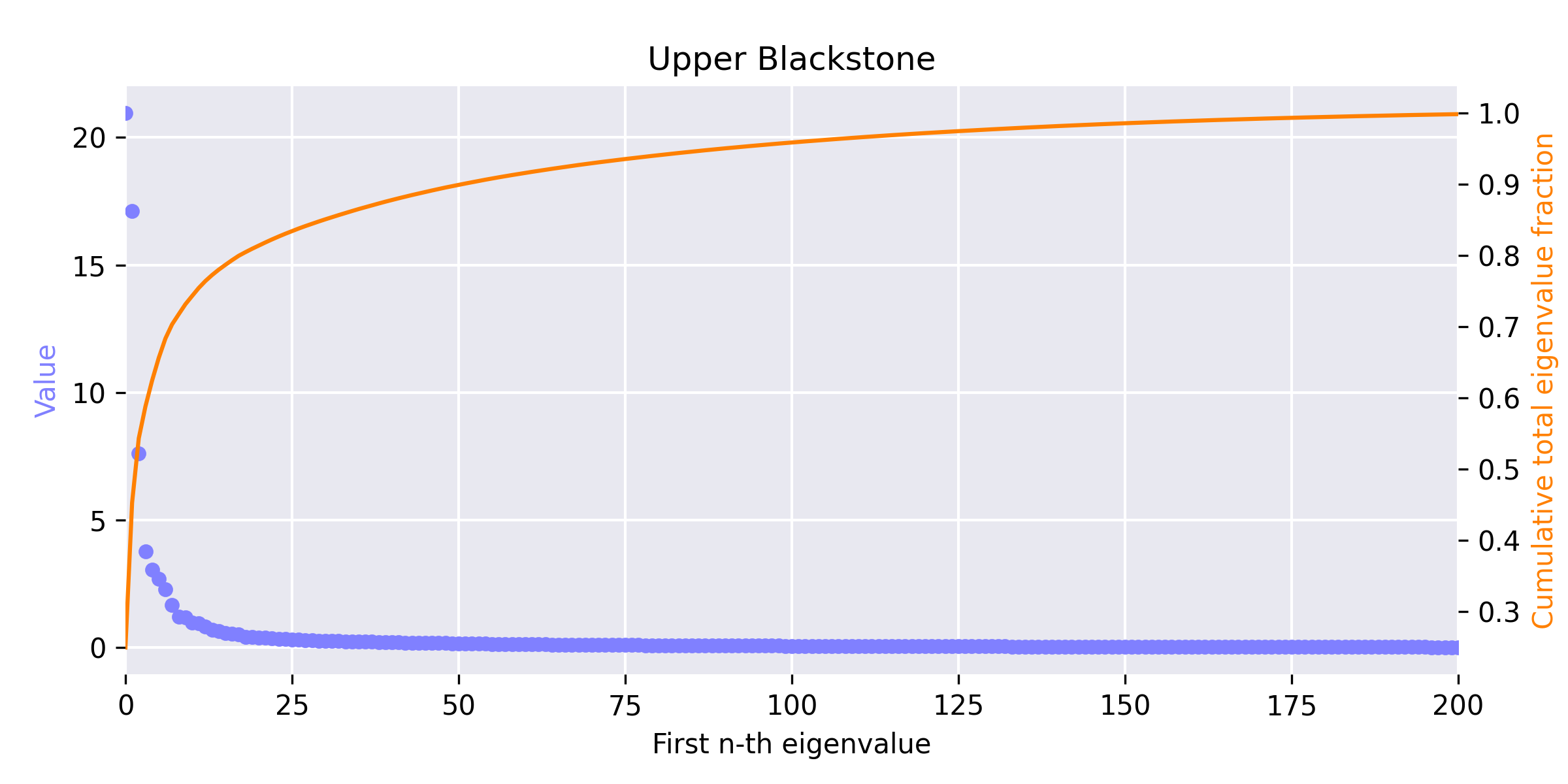

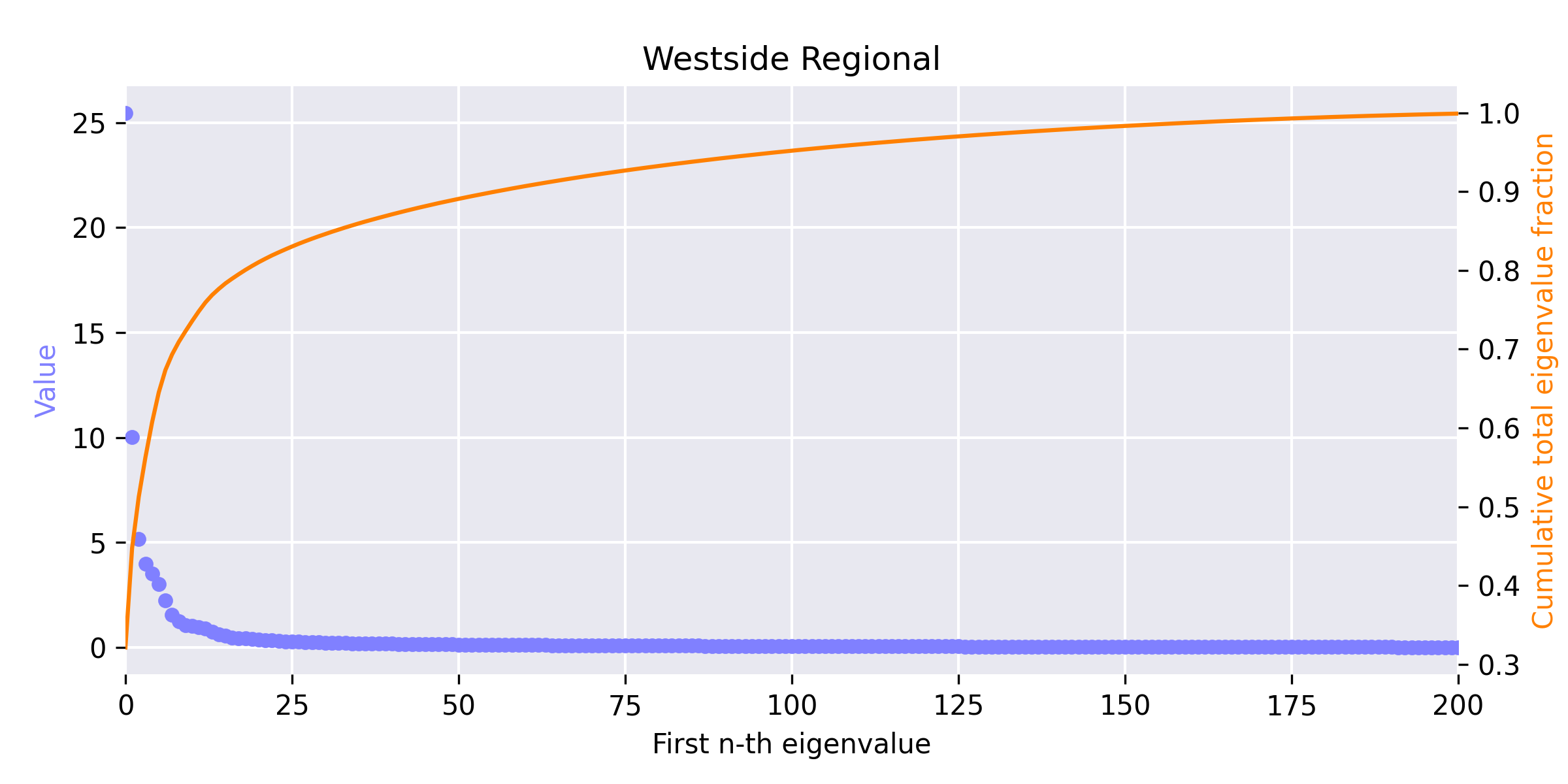

**Figure S4.** Eigenvalue spectra plot for the two larger EBPR SCRS datasets: Upper Blackstone (top, 214 spectra) and Westside Reginal (bottom, 207 samples) selected as representatives for conventional EBPR and S2EBPR categories. The eigenvalues are sorted in descending order and only the first 200 are shown. Neither dataset has a clear “gap-drop” in the eigenvalue profile to identify a group of major eigenvalues. Therefore, we used $p=0.96$ in the kernel divergence calculations.

Brief information, including names, locations, and EBPR configurations of the WRRFs we sampled sludge for SCRS phenotypic survey were presented in **Table S5**. Detailed information of these EBPR facilities can be found in [1-3].

**Table S5.** Supplementary information of source wastewater reuse and reclamation facilities (WRRFs) from which the SCRS datasets were sampled.

| **Facility Name** | **Label** | **Location** | **Configuration** |
| --- | --- | --- | --- |
| Cedar Creek | A | Cedar Creek, KS, USA | S2EBPR, SSM |
| Henderson | B | Henderson, NV, USA | S2EBPR, UMIF |
| South Cary | C | South Cary, NC, USA | S2EBPR, SSR |
| Westside Regional | D | West Kelowna, BC, Canada | S2EBBPR, SSRC |
| Durham | E | Durham, OR, USA | Conventional EBPR, A2O |
| Meriden | F | Meriden, CT, USA | Conventional EBPR, A2O |
| Upper Blackstone | G | Millbury, MA, USA | Conventional EBPR, A2O |
| Westfield | H | Westfield, MA, USA | Conventional EBPR, A/O |

Configuration terminologies and abbreviations:

- EBPR: enhanced biological phosphorus removal
- S2EBPR: side-stream EBPR
- RAS: return activated sludge
- A2O: anaerobic-anoxic-aerobic configuration (a type of conventional EBPR)
- A/O: anaerobic-aerobic configuration (a type of conventional EBPR)
- SSM: side-stream mixed liquor suspended solids (MLSS) fermentation (a type of S2EBPR)
- SSR: side-stream RAS fermentation (a type of S2EBPR)
- SSRC: SSR plus carbon (a type of S2EBPR)
- UMIF: unmixed in-line fermentation (main-stream configuration with enabled S2EBPR-like fermentation conditions)

We also conducted a comparative study with the approach proposed by He et al. (2017). The results were shown in **Table S6**.

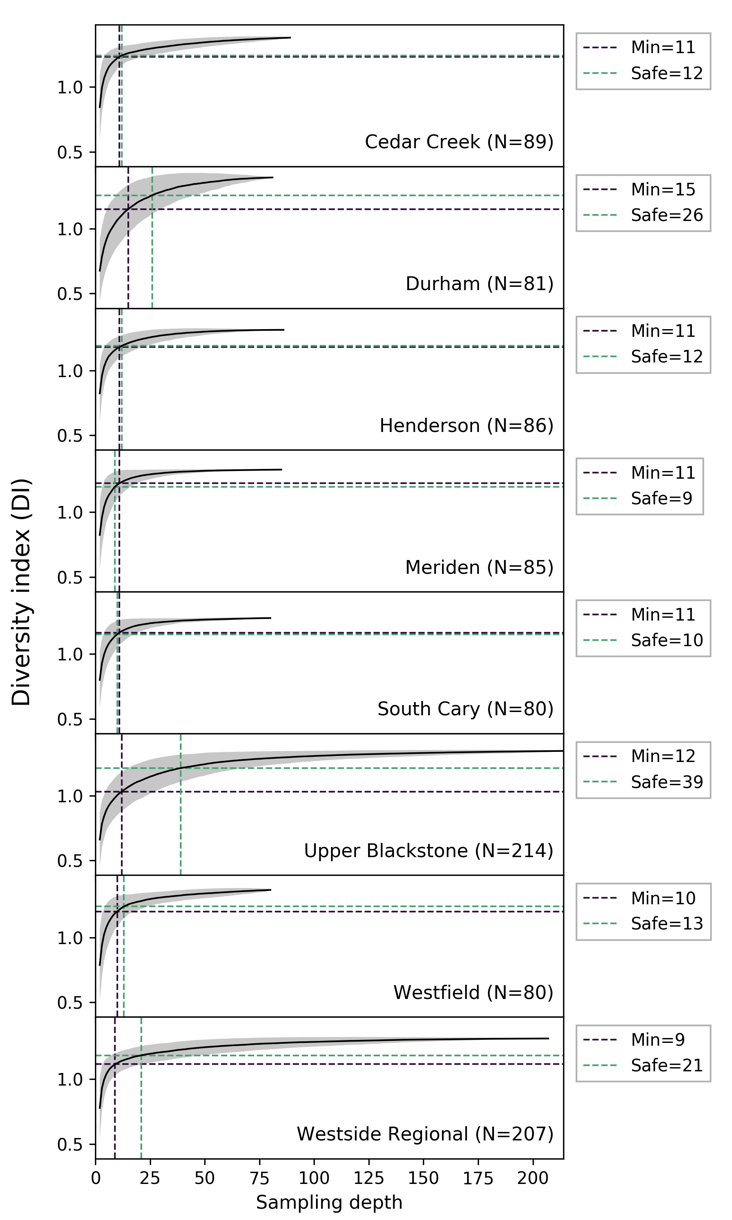

**Figure S6.** Comparative analysis to the minimal sampling depth and safe sampling depth simulated following the approach proposed by He et al, (2017). Solid lines show the DI calculated independently from eight SCRS datasets; averaged from 1000 permutations. The “minimal” sampling depth was determined where DI increment per new sample would be bounded below 0.01, and the “safe” size was where subset DI has reached 90% of dataset maximum. These sampling guidelines were estimated significantly smaller than the kernel divergence approach proposed in this study.
